# Strategies to improve selection compared to selection based on estimated breeding values

**DOI:** 10.1101/2025.01.09.632197

**Authors:** Torsten Pook, Azadeh Hassanpour, Tobias A.M. Niehoff, Mario P.L. Calus

**Affiliations:** Wageningen University & Research, Animal Breeding and Genomics, P.O. Box 338, 6700 AH Wageningen, The Netherlands; University of Goettingen, Department of Animal Sciences, Animal Breeding and Genetics Group, Albrecht-Thaer-Weg 3, 37075, Goettingen, Germany

## Abstract

**Background:** Selection of individuals based on their estimated breeding values aims to maximize response to selection to the next generation in the additive model. However, when the aim is not only about short-term population-wide genetic gain but also the gain over multiple generations, an optimal strategy is not as clear-cut, as the maintenance of genetic diversity may become an important factor. This study provides an extended comparison of existing selection strategies in a unifying testing pipeline using the simulation software MoBPS.

**Results:** Applying a weighting factor on the estimated SNP effects based on the frequency of the beneficial allele resulted in an increase of the long-term genetic gain of 1.6% after 50 generations while reducing inbreeding rates by 16.2% compared to truncation selection based on estimated breeding values. However, this also resulted in short-term losses in genetic gain of 1.2% with the break-even point reached after 25 generations. In contrast, inclusion of the average kinship of an individual to top individuals of the population as an additional trait in the selection index with a weight of 17.5% resulted in no short-term losses and increased long-term genetic gains by 4.3% while reducing inbreeding by 15.8%, achieving very similar efficiency to the use of optimum genetic contribution selection. Combining multiple diversity management strategies, with weights for each strategy optimized using an evolutionary algorithm resulted in a breeding scheme with 5.1% increased genetic gain and 37.3% reduced inbreeding rates. The proposed strategy included the use of optimum genetic contribution, weighting of SNP effects based on allele frequency, average kinship as a trait in the selection index, avoiding matings between related individuals, and lowering the proportion of selected individuals.

**Conclusions:** The combination of selection strategies for the management of genetic diversity was shown to be far superior to the use of any singular method tested in this study. As an efficient use of methods for the management of genetic diversity and inbreeding does not necessarily lead to short-term losses in genetic gain and comes at no extra costs, it is critical for breeding companies to implement such strategies for long-term success.

## Introduction

Although there are substantial differences between livestock and plant breeding species, the general goal of breeding to improve desired traits or characteristics in the breeding organism is universal. Although there might be differences in the details, most breeding programs make use of selection and diversity management with concepts sharing similar ideas across species. To this end, the selection of individuals to be used as parents for the next generation is of critical importance, with the use of estimated breeding values (EBVs) maximizing response to selection.

Although the importance of genetic diversity in a breeding program is a generally accepted fact (e.g., Breeder’s Equation [1]), diversity management in the short-term is usually associated with reduced genetic gains, which for a breeding company can be associated with lower quality short-term product and the associated risk of going out of business when putting too much emphasis on genetic diversity. However, increasing signs of reduced heritabilities [2, 3], increased genetic load [4, 5], and increased expression of inbreeding depression [6] are reported in practice which increases the need for more efficient management of genetic diversity without compromising short-term genetic gain too much.

To assess and compare breeding programs, the use of stochastic simulations provides a powerful tool to generate a digital twin of a breeding program, which can then be used to precisely analyze the impact of specific changes. Compared to the use of deterministic calculations based on quantitative genetic theory [7], this comes with the disadvantages of higher computing costs and calculations only resulting in a realization of a stochastic process instead of an expected value. However, these downsides are compensated by the added flexibility of stochastic simulations – particularly as there is usually no quantitative theory to analyze the exact impact of subtle changes on the outcomes of a breeding program. Software for stochastic simulation includes MoBPS [8], AlphaSimR [9, 10], Adam [11], and QMsim [12].

Concepts for the management of genetic diversity are multifaceted both in complexity and method, with theoretically optimal solutions often being impractical or overly complex to implement. In the following paragraphs, we will provide a general overview of different approaches. The focus here is on optimizing the selection step and selection criteria and explicitly not on restructuring of the breeding program design as a whole to introduce outcross material [13, 14] or splitting the breeding population into subgroups [15, 16].

The gold standard in livestock breeding is the use of optimum genetic contribution selection (OGC) [17] to calculate the contribution of each individual to a subsequent generation to maximize genetic gain under a target inbreeding rate or minimize the inbreeding level under a target genetic gain. Concepts of OGC can be combined with minimum coancestry mating [18, 19] to mate individuals with genetically more distant partners can further reduce inbreeding. In reality, an implementation of these concepts in their entirety can however be difficult. For example, it is usually not possible to have a single offspring from a sow as a litter contains multiple individuals, resulting in more or less offspring than theoretically wanted. Similary, in plant breeding, logistical limitations and field design can lead to required simplifications. Simplifications might include selecting a maximum number of individuals from a given family or avoiding matings between close relatives.

Over the years a variety of alternatives to OGC have been proposed with various objectives and/or motivations behind them. For instance, Goddard [20] suggested putting more emphasis on rare alleles, as they are more likely to get lost due to drift, hitchhiking or unfavorable correlation with closely linked other quantitative trait loci (QTLs). Building upon the work of Goddard [20], Jannink [21] suggested a simplified weighting factor of 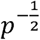 with a simulation study by Liu et al. [22] demonstrating the long-term benefits of the approach.

As selection of individuals in a breeding program is typically done based on a variety of traits that are combined in a selection index, another alternative is to include the contribution of an individual to the genetic diversity of the population as a pseudo-trait in a selection index. An example of this can be found in the Net Merit index used in US dairy cattle breeding [23].

In addition to the expected genomic value of a hypothetical offspring (“breeding value”) the variance among breeding values of offspring can be considered in selection decisions. Hence, an individual with a lower expected genomic value can still be of value when the goal is to generate some high-performance offspring individual. The variation in inherited gametes of a potential parent, which reflects the Mendelian sampling variance (MSV), plays a key role in this context. Several approaches to integrate MSV into breeding strategies have been proposed with the primary focus on short-term gain genetic [24–29]. However, some studies also highlight the long-term benefits of MSV integration, such as enhanced genetic gain and diversity with Allier et al. [30, 31] specifically targeting long-term “usefulness” of individuals in breeding decisions.

Lastly, approaches such as MateSel [32] and AlphaMate [33] can be used to generate a specific mating plan. A potential benefit from this is that it is much easier to implement restrictions or constraints to the selection of individuals, e.g. to select a certain number of individuals which will all have the same number of litters, compared to the frequentist / Lagrange-based procedure of OGC which just dictates the optimal contribution of all considered individuals [17].

As most studies compare their newly developed approaches exclusively against plain selection based on EBVs rather than against other selection strategies for the maintenance of genetic diversity, the goal of this paper is to compare different approaches with each other. Furthermore, an evolutionary optimization pipeline [34] is employed to assess whether and how approaches can be combined and provide suggestions for which methods to use in real-world breeding programs. A further aim is to provide a uniform testing pipeline for breeders and researchers to easily compare newly developed approaches with common standards in the field as well as approaches tested in this manuscript.

## Methods

As a baseline breeding program for optimization, we consider a generic breeding program simulated using the R-package MoBPS [8]. In brief, a genome with 10 chromosomes of 2.5 Morgan length and 25,000 equidistantly spread SNPs was simulated. A single quantitative trait with 1,000 purely additive QTLs and a heritability of 0.3 was simulated, with the QTLs being a subset of the SNPs. We are here considering 50 generations of breeding with truncated selection, in which the top 40 males and top 100 females are used as sires and dams for the breeding nucleus of the next generations. The selected individuals have equal contributions to the next generation. All individuals were genotyped and phenotyped before selection based on a genomic best linear unbiased prediction (GBLUP) [35, 36] that includes individuals from the current generation.

To initialize the population with a realistic population structure, 1,000 founder individuals with randomly sampled genotypes were generated and subsequently 10 generations of mating with phenotypic selection as a burn-in phase were simulated. Each subsequently proposed scenario was simulated 50 times (unless specifically indicated otherwise), with reported values representing averages across replicates.

Subsequently, we are considering various modifications to the breeding program that will be discussed in the following subsections. To mimic the generation of production individuals, an additional cohort of production individuals was generated with sires and dams selected based on their EBVs – these individuals were not used for subsequent selection but only for evaluating the output of the breeding program. Genetic gains are evaluated based on the underlying true genomic values of the production individuals based on the underlying known QTL effects. The genetic variance is computed based on the underlying true genomic values and the inbreeding levels are derived based on Identify-by-descent [37] as the originating founder for each chromosome segment is tracked in simulation [8].

### Weighting allele effects by allele frequency

Following the methods suggested by Goddard [20] and Jannink [21], we are considering the use of a modified selection criterion in which marker effects are weighted based on the allele frequency of the beneficial allele in the population:

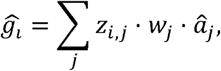

With u·_*j*_ being the estimated effect of marker j, z_*i,j*_ being the genotype of individual i at marker j and w_*j*_ being the weighting factor of marker j. The weighting factors are calculated based on the allele frequency of the beneficial allele p_*j*_, determined based on the sign of u·_*j*_. We are here considering the following potential weightings:

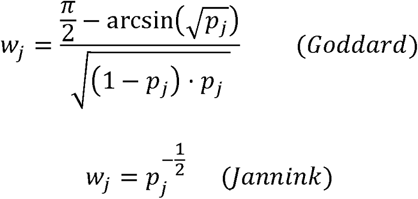

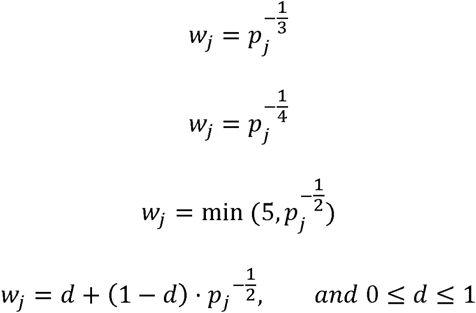

With the first two weighting factors being the respective suggestions by Goddard [20] and Jannink [21] while subsequent variants provide more moderate weighting factors to mitigate short-term losses (see Figure 1).

**Figure 1:**
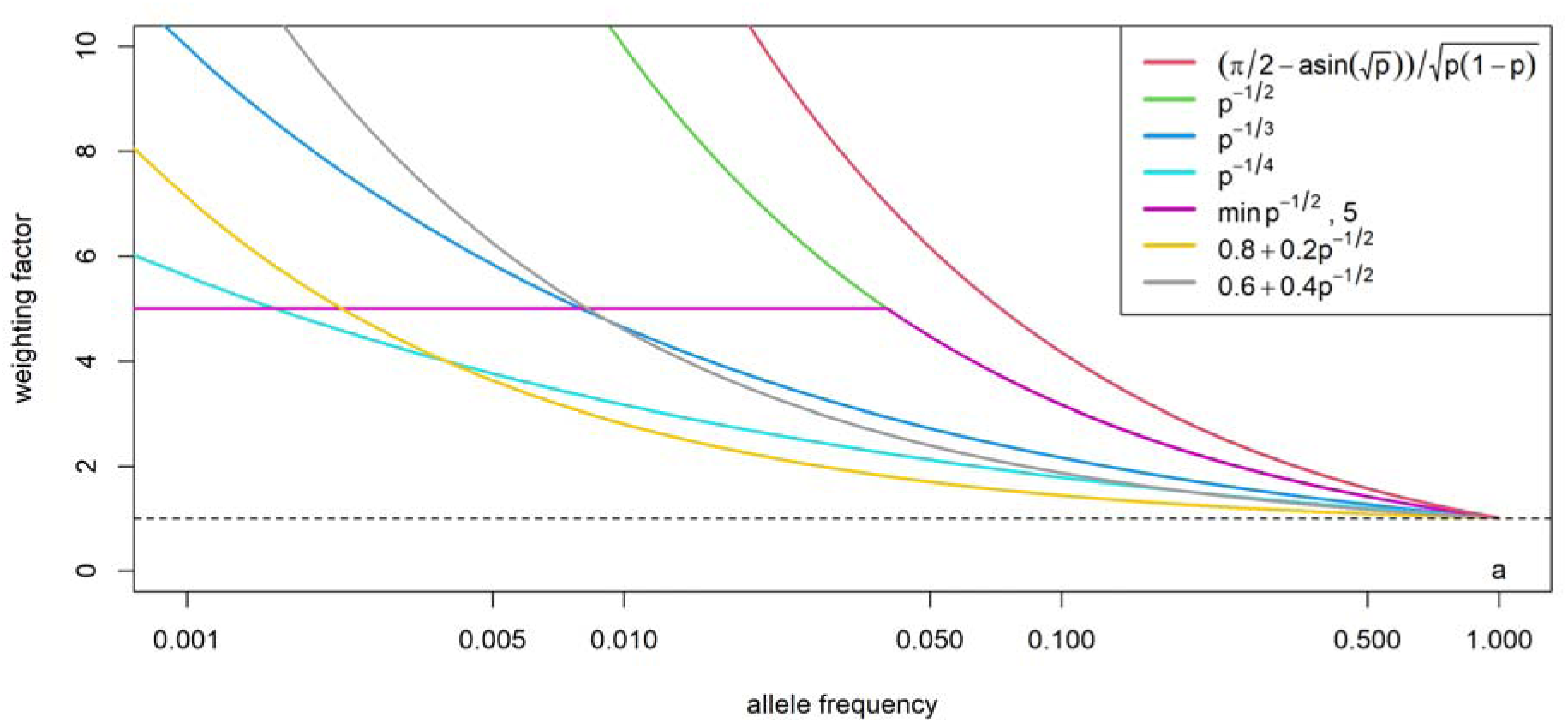
Impact of allele frequency weighting functions. Weighting factor for the different weighting functions depending on the allele frequency of the beneficial allele.

#### Relatedness to breeding nucleus

Similar to the concepts used in the Net Merit index [23], the relatedness of individuals to other individuals in the breeding nucleus can be accounted for as an additional trait in the selection index:

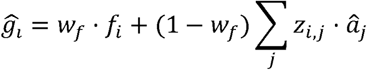

With w_*f*_ being a weighting factor. For 1_*i*_ we are here considering two options:

1. Average kinship to the current breeding population
2. Average kinship to the top individuals

Here, we are considering those individuals that would be selected based on their EBVs as the top individuals. The calculation of w_*f*_ is done by first scaling both the estimated breeding value and *f* to have mean 0 and variance 1 across selection candidates and subsequently weighting these two accordingly with relative weightings between 5 to 40% being considered in the subsequent simulations. Similar considerations can be applied in a multi-trait case by effectively using *f* as an additional trait in a selection index. Pairwise kinships are defined, as the probability of two randomly chosen alleles from two individuals at the same locus to be identical by descent [38] and calculated based on pedigree with a pedigree-depth of 7.

#### Avoiding matings between related individuals

We are here considering different extents of mating control. Firstly, we consider avoiding matings between full and/or halfsibs as such matings would naturally lead to high inbreeding levels in their offspring. To not only exclude these extreme cases, we propose a more advanced selection strategy that aims at avoiding matings between individuals that would lead to a high inbreeding level in general. For this, the expected inbreeding of a mating pair is calculated using on their pedigree-based kinship and combinations above a certain threshold are avoided. As a threshold, we are proposing to use a quantile of the resulting expected inbreeding levels from all mating combinations. We are here considering values between the 10 to 90% quantiles – e.g. for the 90% quantile in a population with 40 sires and 100 dams of the potential 40,000 matings only 36,000 sire-dam combinations would be potential matings. Note that contributions of sires and dams to the next generation can change if an individual has high expected inbreeding rates to a particularly high/small share of individuals of the other sex as finally executed matings are sampled from the set of all allowed matings. Overall differences in contributions of individuals should be small. Finally, instead of expected inbreeding, the expected kinship of a potential offspring to the current breeding population was used to avoid matings. Again, quantiles are used as thresholds. As expected kinship is the mean of the kinships of the parents to the population, this will heavily reduce contributions of sires and dams with high average kinship to the current breeding population and thereby potentially even result in some of the selected individuals not being used as sires or dam.

#### Optimum genetic contribution

For the use of optimum genetic contribution [17], the R-package optiSel [39] is used to calculate individual contributions. To mimic the common restriction of animal breeding programs that contributions on the female side cannot be easily controlled, optimum genetic contribution was only applied on the male side [39]. For this, optiSel is provided with information on all males and the already selected females with the goal of maximize genetic progress (target = “max.BV”, uniform = “female”) under the restriction of an increase of avg. kinship to the reference population with thresholds tested between 0.3% and 0.7%.

#### Mate allocation

For mate allocation algorithms, the software AlphaMate [33] was considered. AlphaMate was used to suggest 1000 matings from 40 males and 100 females to use same selection proportions as the baseline scenario with equalized contributions of the selected individuals (EqualizeMaleContributions = Yes, EqualizeFemaleContributions = Yes). To balance genetic gain and genetic diversity the angle parameter in AlphaMate was used with values ranging from 5 to 45 degrees with an angle of 0 corresponding to only selection for genetic gain and an angle of 90 corresponding to only selecting for genetic diversity. To reduce computing time, the convergence criteria and the maximum number of iterations performed in AlphaMate were reduced (nEvolAlgNumberOfIterations = 1000; nEvolAlgNumberOfIterationsStop = 50).

#### Mendelian Sampling Variance

To assess the potential of accounting for MSV, we considered two approaches: Firstly, Bijma et al. [25] suggested theoretically motivated indices to select based on the probability of hypothetical offspring of an individual to be selected (*I*_*5*_, [25]) and to maximize response to recurrent selection (*I*_*6*_, [25]). Secondly, we are considering the ExpBVSelGrOff criterion given in Niehoff et al. [29] that is motivated by maximizing genetic response in the grand offspring without pre-selection and accessed based on the full breeding nucleus. As prediction accuracies for MSV were low, we additional considered the inclusion of the last six generations of individuals in the estimation.

#### Adapted selection proportions

Lastly, we are considering a modification of the selection proportions. In the baseline, the number of selected sires is 40% of the number of selected dams, meaning that we are here only considering the same relative changes for both sexes and considering the selection of between 28 and 50 sires and 70 and 125 dams. Although equally correct when this is the only modification of the selection strategy, we avoid the term “selection intensity” throughout the manuscript, as when combined with other selection strategies, the selection of less individuals will not necessarily increase the selection intensity when individuals with lower EBVs are selected.

#### Joint optimization (Evolutionary Algorithm)

To perform an optimization of the breeding program under consideration of multiple of the previously suggested approaches, we are using here an evolutionary algorithm based on the pipeline suggested by Hassanpour et al. [34].

For this, we first define a target function that includes short- and long-term genetic gain and remaining genetic diversity. For genetic gain, we are considering the average underlying true genomic value in the breeding population *g*_*t*_ with equal weights on each year *t*. For genetic diversity, we are considering the resulting inbreeding level *f*_*50*_ and the genetic variance 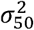 after 50 generations. The three components are first scaled by 25, 0.045, and 0.3, reflecting the standard deviations across different scenarios tested. Subsequently, genetic gain was weighted with a factor of 2/3 compared to 1/6 for both genetic diversity metrics:

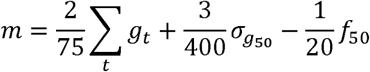

For the optimization problem, we are considering the nine parameters subject to optimization, as given in Table 1. This includes four parameters to account for relatedness to top individuals (*f*_i,1_) , the full population (*f*_i,2_), the weighting of estimated SNP effects based on Goddard [20], and a more moderate weighting of 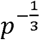 resulting in the following formula for the selection criterion:

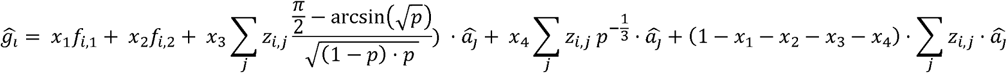

**Table 1:**
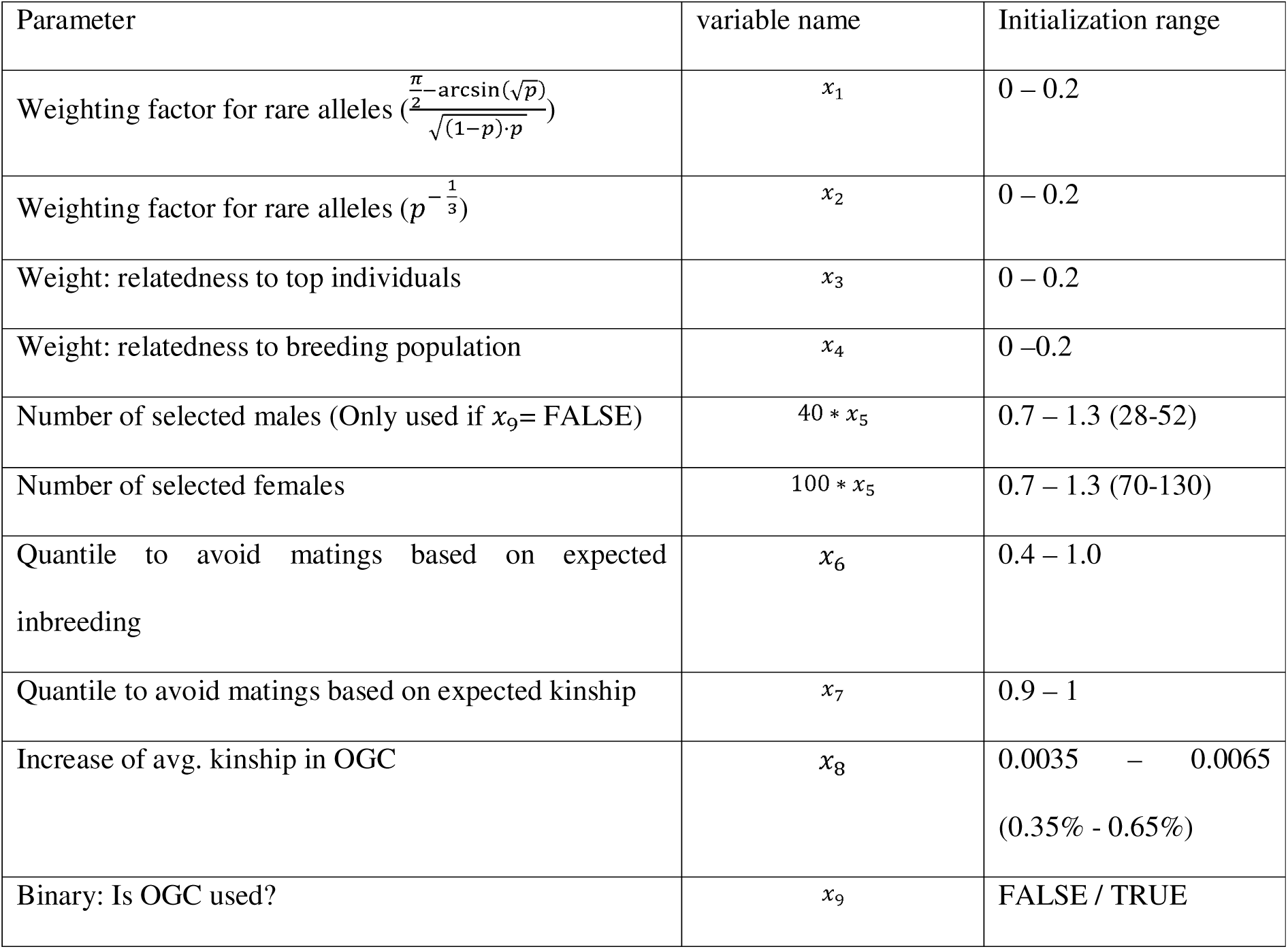
Overview of parameters considered in the evolutionary pipeline.

Furthermore, avoiding matings based on expected inbreeding and kinship are considered as two additional parameters, while the selection proportion on the male and female side is treated as a joint parameter with the aim to select 60% less males than females. Lastly, a binary parameter to control if OGC is used on the male side is used and if so a further parameter to control the constraint of inbreeding rate in OGC is used. Note that if OGC is used, selection proportion on male and female side can deviate from the 40:100 ratio.

Initialization for the binary parameter done by random sampling from a Bernoulli random variable with probability of 0.5, while all other parameters are sampled from a uniform distribution with initialization ranges given in Table 1. No constraints for budget or similar are applied that would require scaling or a more complex initialization.

As a second scenario for the use of the evolutionary algorithm, we consider the use of a modified target function that places more emphasis on short-term genetic gain. For this, the weighting of genetic gain is multiplied by an interest factor ( i = 1.03). Additionally, the weighting of genetic gain was increased by a factor of 2.5 to offset the lower variation in the genetic gain term due to the interest scaling:

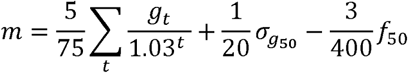

As the evolutionary pipeline by Hassanpour et al. [34] does not reach convergence per se, the final optima were determined based on visual inspection of the suggested optima from the last couple of iterations. The determined breeding schemes were subsequently simulated 50 times.

## Results

In the baseline scenario a genetic gain of 26.19 genetic standard deviation (gSD) was obtained after 50 generations of breeding. With regard to genetic diversity, inbreeding levels increased from 0.056 in generation 1 to 0.680 in generation 50 with an inbreeding rate of about 1.5% in the first couple of generations, while the standard deviation of underlying true genomic values was reduced to 28.9% of the initial population with 64.0% of all SNPs and 74.5% of all QTLs (77.1% of those for the favorable variant) being fixed (Figure 2). As different metrics for genetic diversity resulted in very similar rankings between scenarios, we will focus on genetic gain and inbreeding in the following description of results. The interested reader is referred to Supplementary File S1 - S6 for a detailed per generation overview on genetic gain, remaining genetic variance, and share of QTLs fixed for all considered scenarios.

**Figure 2:**
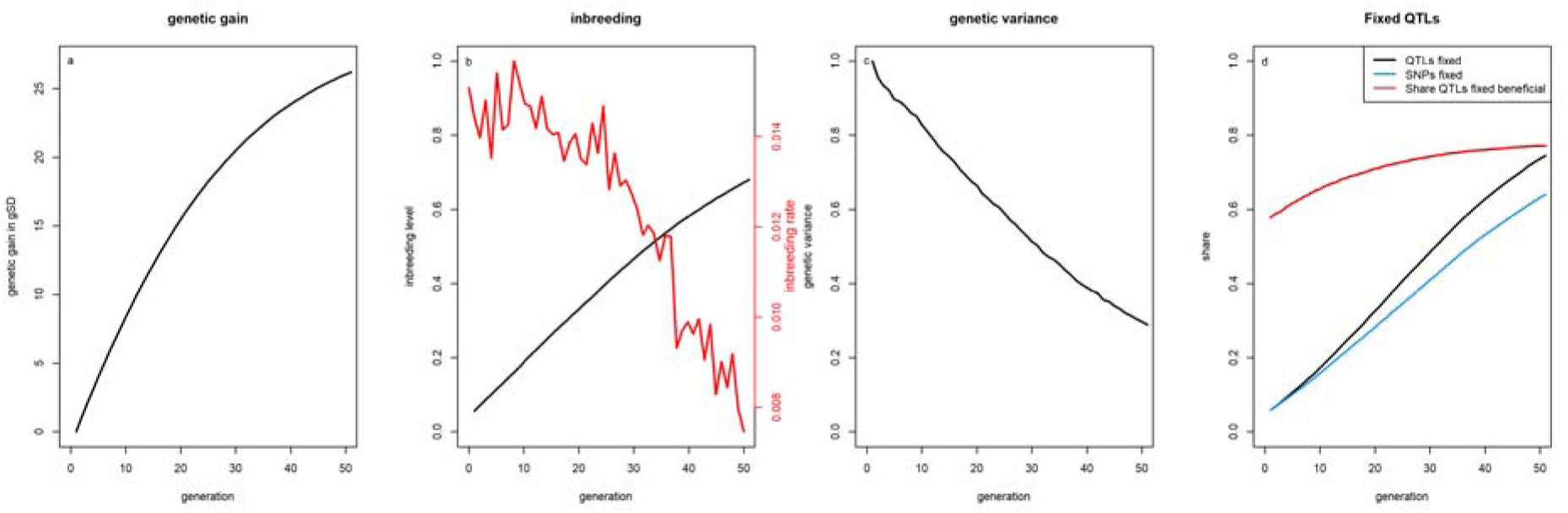
Genetic gains, inbreeding levels, genetic variance and share of fixed QTLs in the reference scenario. Genetic gains (a), inbreeding levels and rates (b), genetic variance (c), and share of SNPs and QTLs fixated (d) for the baseline scenario with selection based on estimated breeding values.

### Weighting allele effects by allele frequency

With the theoretical optimal weightings [20], highest long-term genetic gains with a relative improvement of 0.41 gSD (+1.6%) compared to the baseline were obtained, with a reduction of the inbreeding level after 50 generations of 0.102 (-16.2%) (Figure 3). However, there are short-term losses in genetic gain of up to 0.15 gSD (-1.2%) and a break-even point for genetic gain is only reached after 25 generations. Although not as beneficial in the long-term, applying a weaker relative weighting on rare alleles 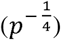 resulted in almost no short-term losses (-0.01 gSD; -0.1%) while maintaining about half of the long-term (+0.23 gSD; + 0.9%) and still substantially reduced inbreeding rates (-0.037; -6.0%). Similar results were also obtained when assuming SNP effects to be known (Supplementary Figure S7).

**Figure 3:**
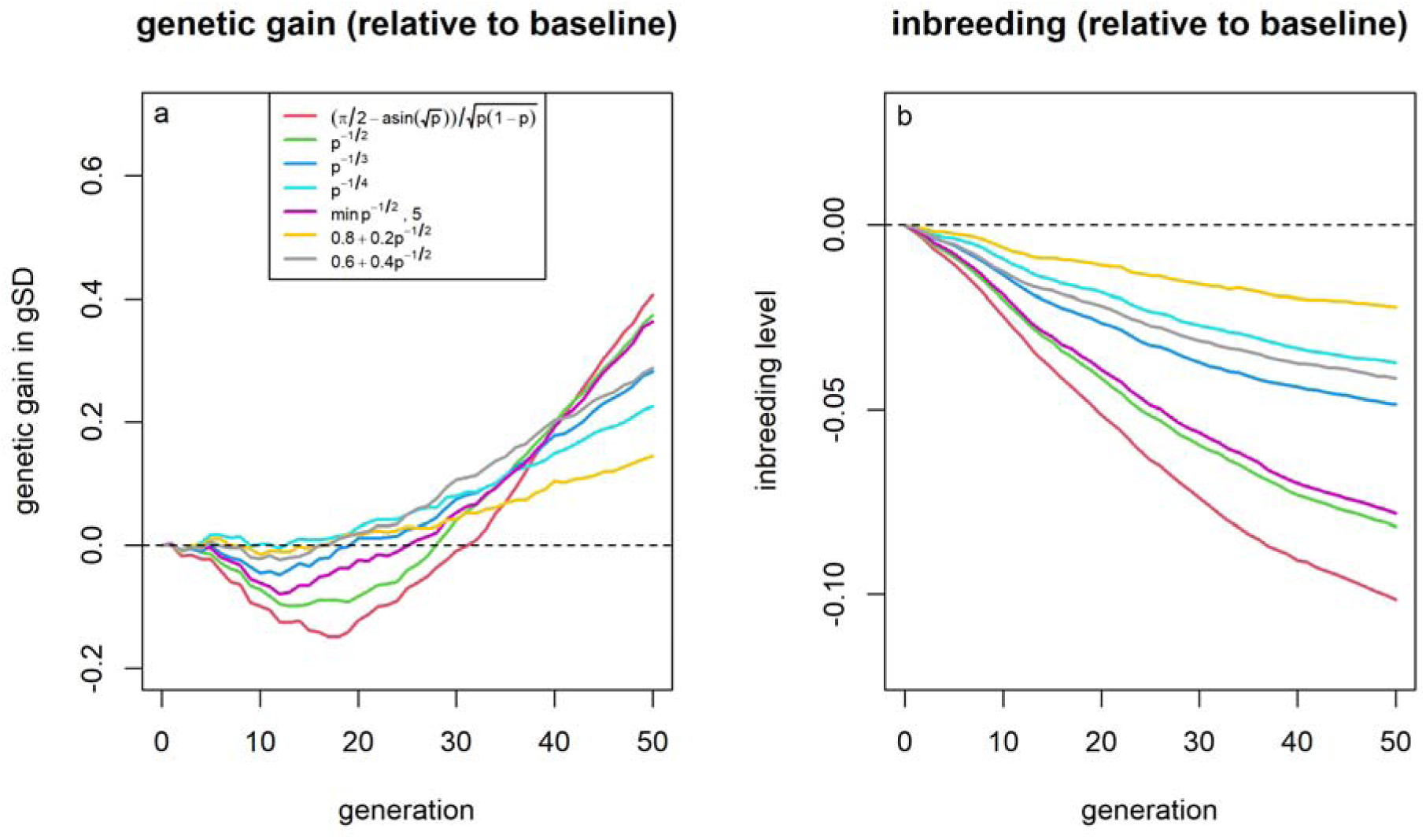
Genetic gain and inbreeding levels in the allele frequency based effect weighting scenarios. Genetic gains (a) and inbreeding levels (b) relative to selection based on estimated breeding values for different weighting factors for SNP effects depending on the allele frequency of the beneficial variant. Due to the small overall impact of allele weighting strategies and high variance in outcomes, figures were generated based on 500 replicates to reduce variance.

#### Relatedness to breeding nucleus

As long as the relative weight for the average kinship of an individual to the top individuals in the current breeding cycle is below 20%, basically no short-term losses were observed with a weighting of 17.5% leading to +1.11 gSD (+4.3%) and 0.099 (-15.8%) lower inbreeding levels after 50 generations (Figure 4). Relative to this, selection based on the average kinship of an individual to the overall population was less efficient with lower genetic gains and a weaker reduction in inbreeding levels (see Supplementary Figure S8)

**Figure 4:**
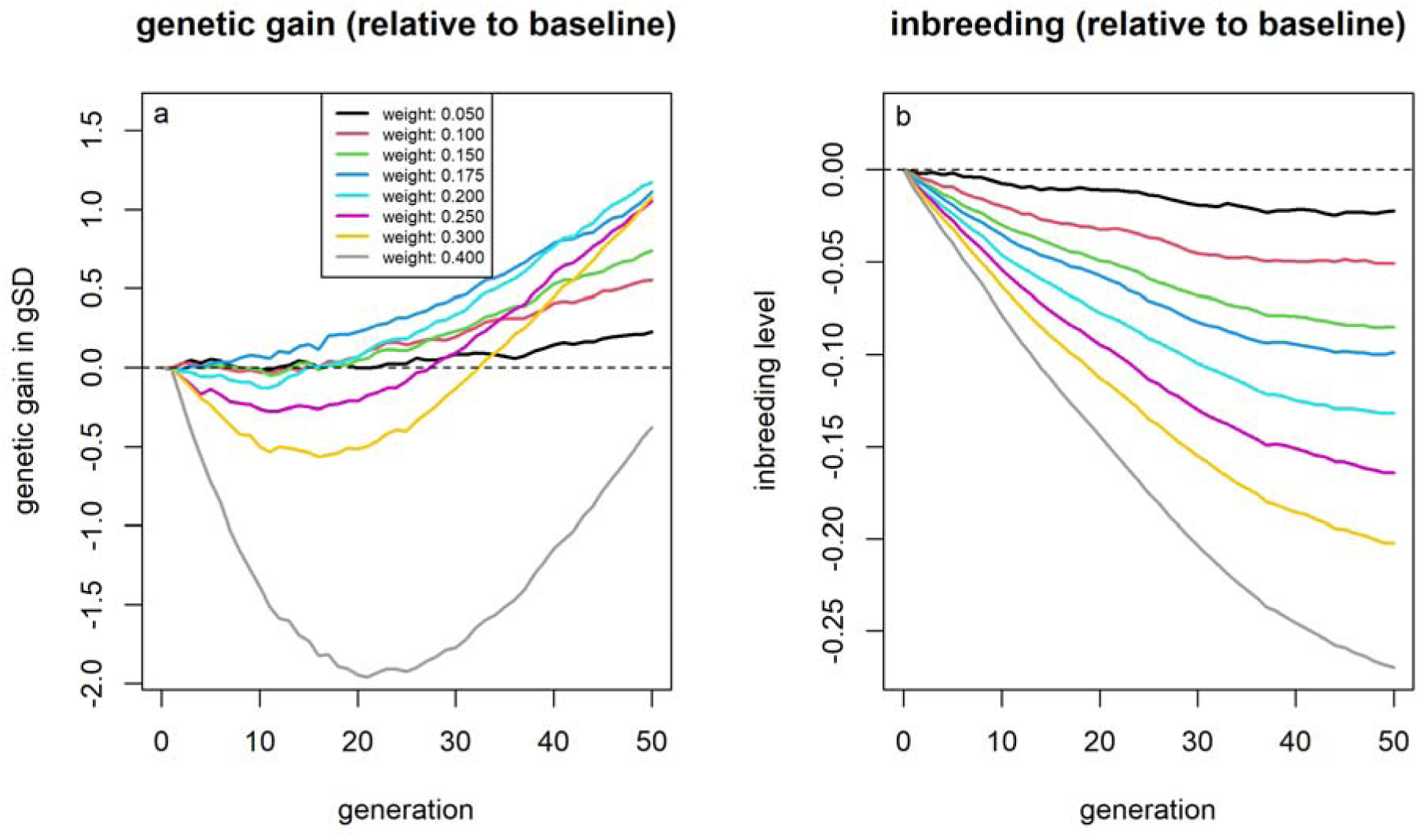
Genetic gain and inbreeding levels of the scenarios including average kinship in the selection index. Genetic gains (a) and inbreeding levels (b) relative to selection based on estimated breeding values for different index weights for the average kinship of individual to the individuals with the highest EBVs from the current population.

#### Avoiding matings between related individuals

Avoiding fullsib matings had no noticeable effect on both genetic gains and inbreeding as the number of such cases is quite limited. When also avoiding halfsib matings, inbreeding levels were noticeably reduced (- 0.045 / -7.3%) with slightly higher genetic gains (+0.27 gSD; +1.0%) after 50 generations (Figure 5).

**Figure 5:**
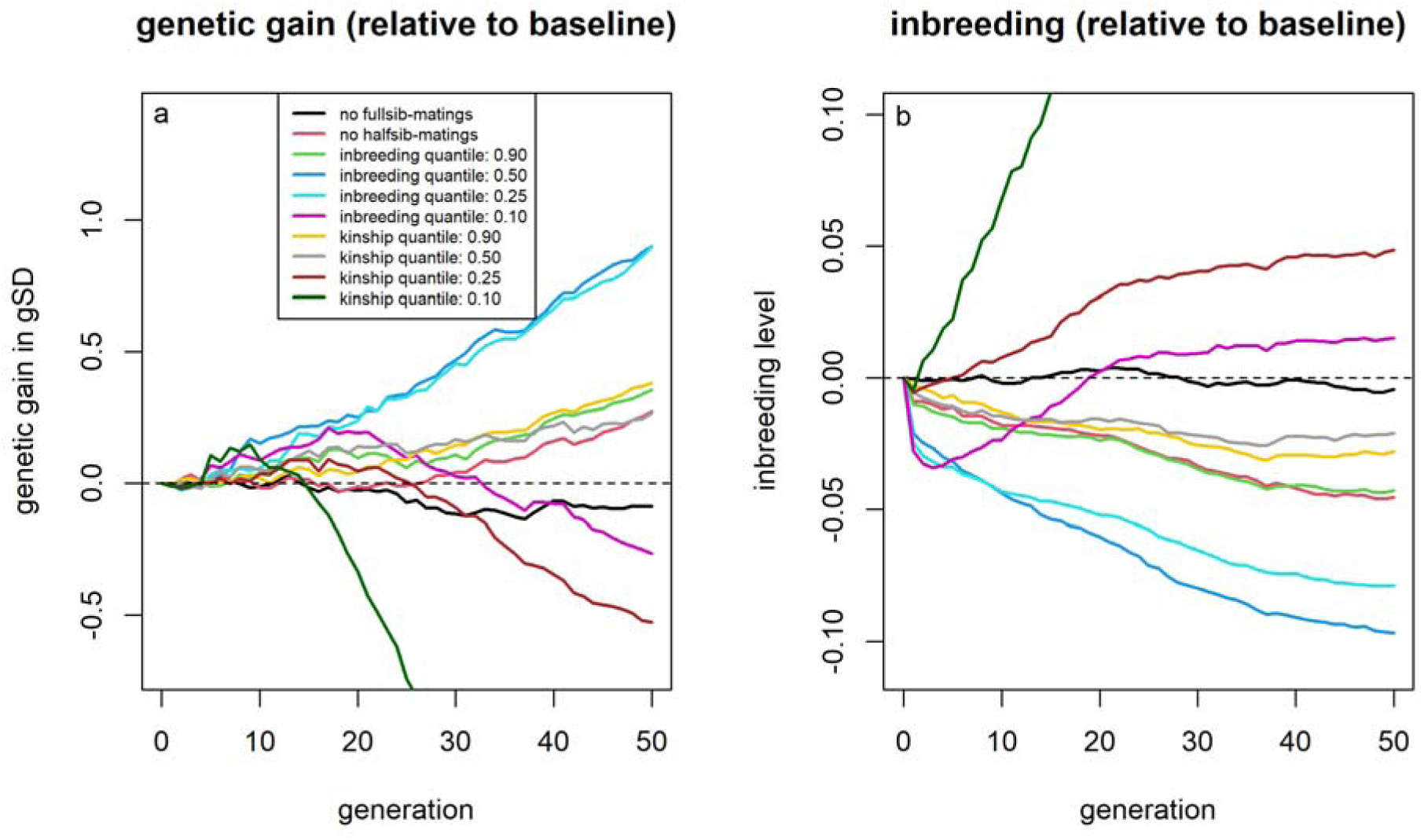
Genetic gain and inbreeding levels of the scenarios including avoiding matings between specific sire-dam combinations. Genetic gains (a) and inbreeding levels (b) relative to selection based on estimated breeding values for strategies to avoid matings between specific sire-dam combinations.

Avoiding matings between individuals that would result in high expected inbreeding levels showed even higher improvements with use of the 50%-quantile as a threshold resulting in an additional genetic gain of 0.90 gSD (+3.4%) and reduced inbreeding levels of 0.097 (-15.5%). Avoiding matings based on expected kinship to the current breeding population reduce inbreeding, however, with a higher share of excluded mating, the higher usage of some individuals as sires/dams led to increases in inbreeding and reduced genetic gains. A detailed overview of the performance with different quantiles for both approaches can be found in Supplementary Figures S9 and S10.

#### Optimum genetic contribution

The use of OGC showed a strict improvement compared to selection based on EBVs, with a constraint limiting the increase in inbreeding to at most 0.5% based on selection on the male side, leading to higher short- and long-term genetic gain (+0.59 gSD; + 2.3%) and reduced inbreeding (-0.062; -10.0%). Stronger restrictions on the increase in inbreeding lead to even higher long-term genetic gain and reduced inbreeding but negatively impact short-term genetic gain (Figure 6).

**Figure 6:**
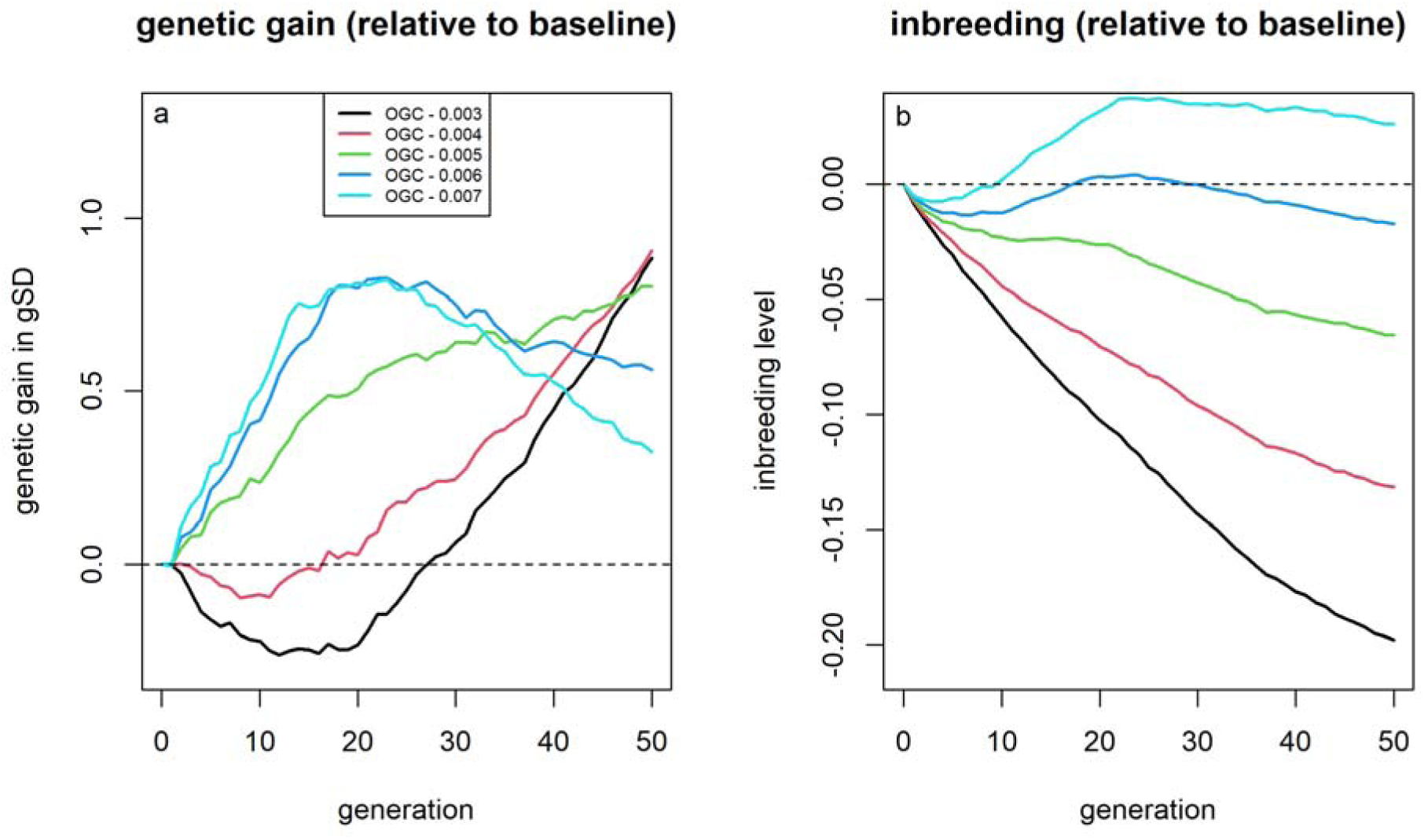
Genetic gain and inbreeding levels of the scenarios using optimum genetic contribution (OGC). Genetic gains (a) and inbreeding levels (b) relative to selection based on estimated breeding values for different inbreeding constraints when using optimum genetic contribution selection.

#### Mate allocation

Using AlphaMate led to substantially increased long-term genetic gain (+1.17 gSD; +4.5%) and reduced inbreeding (-0.163, -26.2%) when using an angle of 30. Short-term genetic gains were reduced by up to 0.15 gSD (-1.8%) and the break-even point for genetic gain was reached after 25 generations. Increasing the weight on genetic gain (angle = 15) reduced long-term benefits (+0.80 gSD; -0.073 inbreeding level) but avoided any short-term losses (Figure 7). Further increasing the weight on genetic gain (angle = 5) did not result in additional improvements in genetic gain or inbreeding levels.

**Figure 7:**
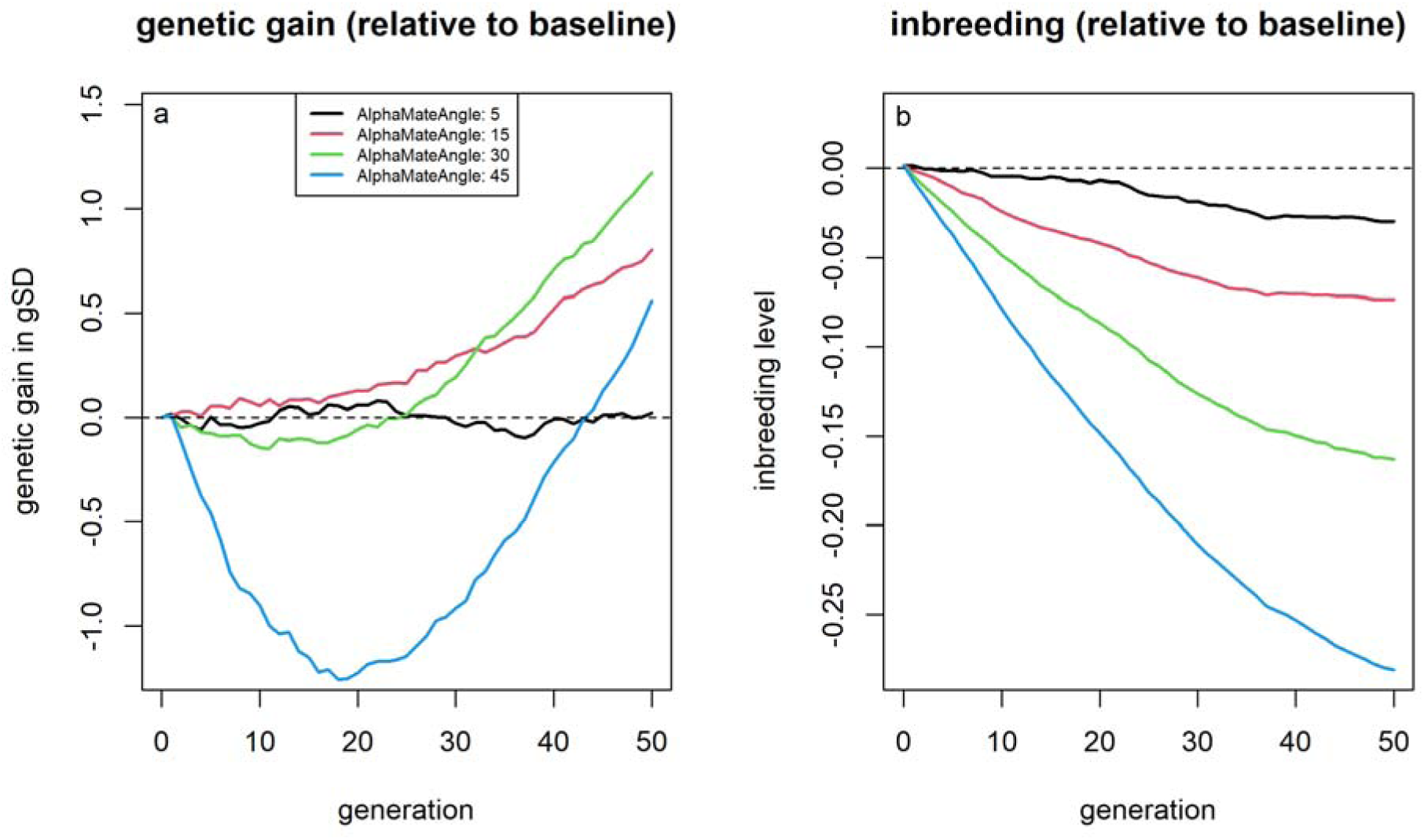
Genetic gain and inbreeding levels of the scenarios using AlphaMate. Genetic gains (a) and inbreeding levels (b) relative to selection based on estimated breeding values for different weightings (angles) in AlphaMate between genetic gain and diveristy.

#### Mendelian sampling variance

All three approaches considered for integrating MSV into breeding led to slightly reduced genetic gain and slightly increased inbreeding levels, compared to selection based on estimated breeding values (Figure 8). Prediction accuracies for MSV were on average 0.17 across the 50 generations compared to an average accuracy of 0.53 for the EBVs. When including six generations in the breeding value estimation accuracies increased to 0.32 and 0.58 respectively with genetic gains and inbreeding levels on par with the baseline scenario (Supplementary Figure S11).

**Figure 8:**
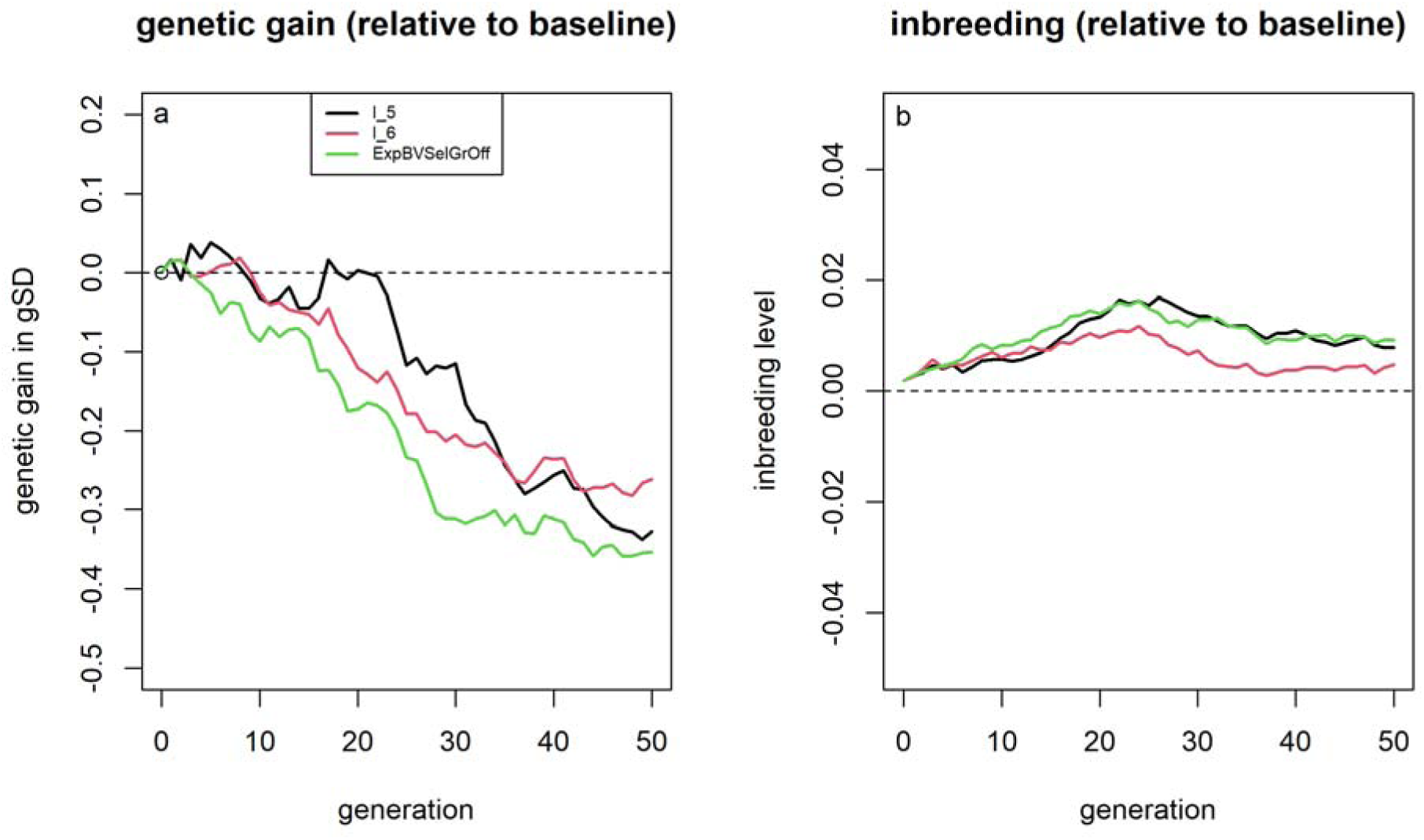
Genetic gain and inbreeding levels of the scenarios accounting for Mendelian sampling variance in selection. Genetic gains (a) and inbreeding levels (b) relative to selection based on estimated breeding values for different approaches to account for Mendelian sampling variance in selection.

#### Adapted selection proportions

By decreasing the proportion of selected individuals, short-term genetic gains can be substantially increased with a 30% decrease of selection proportions resulting in up to 1.01 gSD (+10.0%) higher genetic gain (Figure 9). However, this increased inbreeding rates by 42%. While selecting less individuals led to higher gains for the first 20 generations of breeding, genetic gains subsequently reduced with the break-even point reached between 35 and 50 generations. On the other hand, a higher proportion of selected individuals yielded smaller short-term gains, but became slightly beneficial in regard to genetic gain after 50 generations. Increasing short-term genetic gain by 1% by modifying selection proportions resulted in an increase of inbreeding rates of 4 to 5% across scenarios.

**Figure 9:**
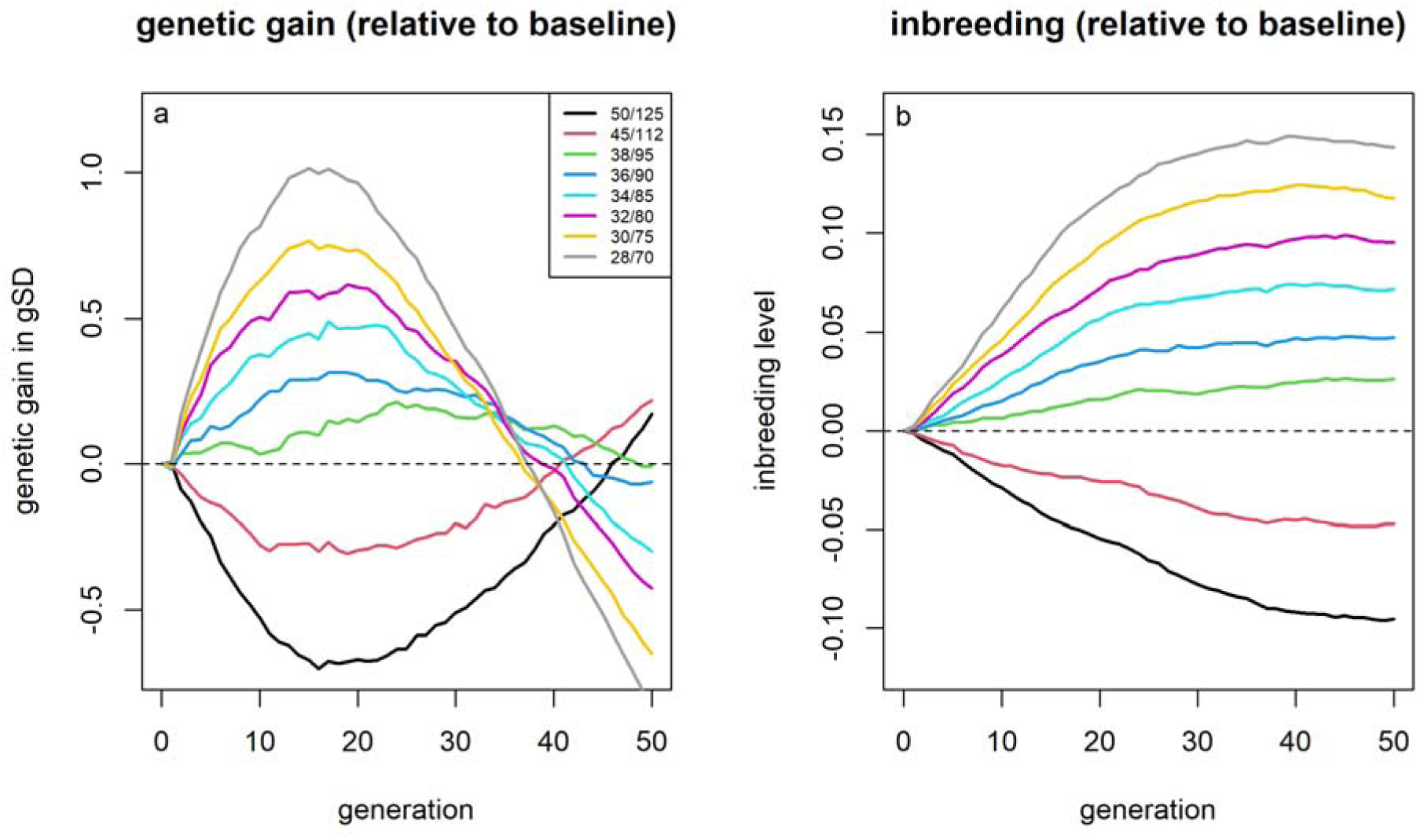
Genetic gain and inbreeding levels of the scenarios with different selection proportions. Genetic gains (a) and inbreeding levels (b) relative to selection based on estimated breeding values depending on the used selection proportion.

#### Joint optimization

The use of the evolutionary pipeline for both target functions suggests the joint use of various previously analyzed selection methods for the optimal management of genetic diversity and genetic gain (Table 2). Note that the evolutionary pipeline does not suggest an optimum per se, which makes the choice of the finally chosen optimum somewhat arbitrary and open to interpretation, however in both cases no major improvements of the target function after iteration 50 of the pipeline were obtained. The interested reader is referred to Supplementary Figures S12 and S13 for the respective suggested optima for each iteration of the evolutionary pipeline and their estimated values for the respective target functions.

**Table 2:**
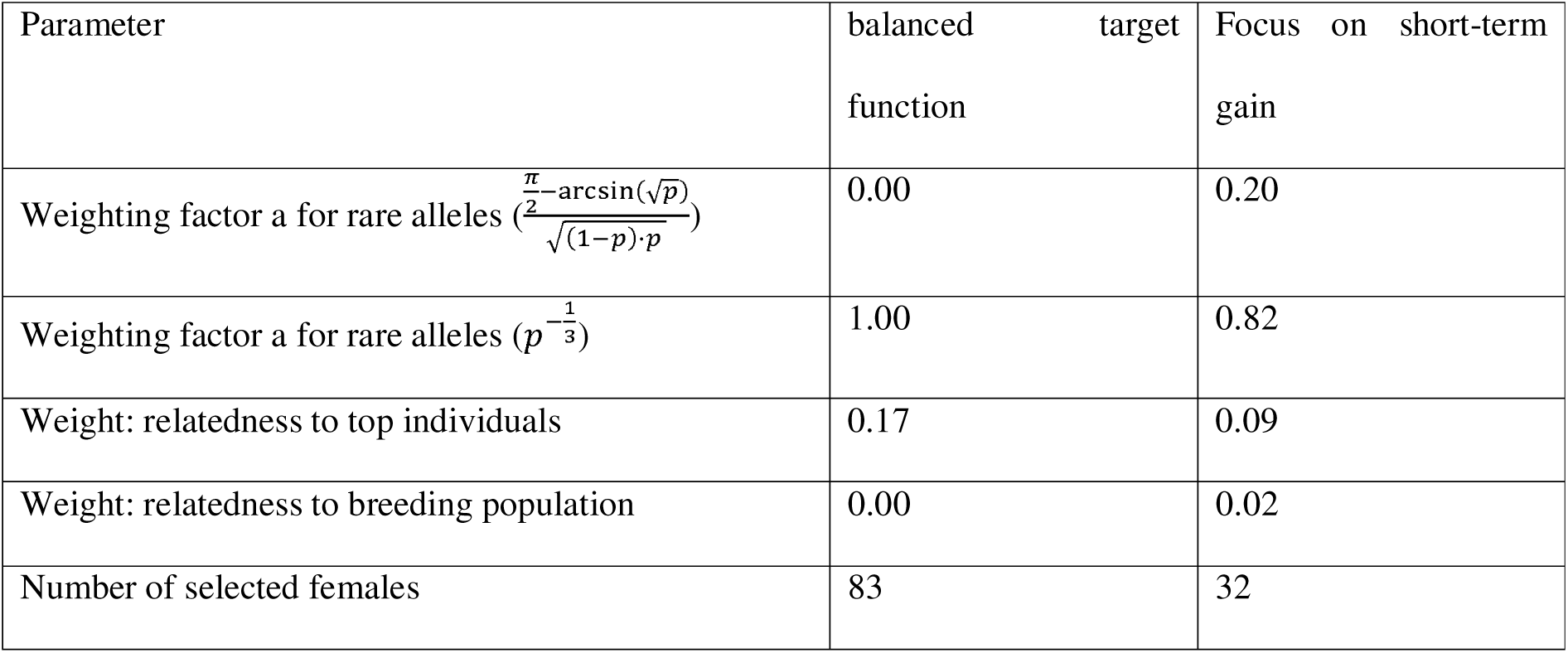
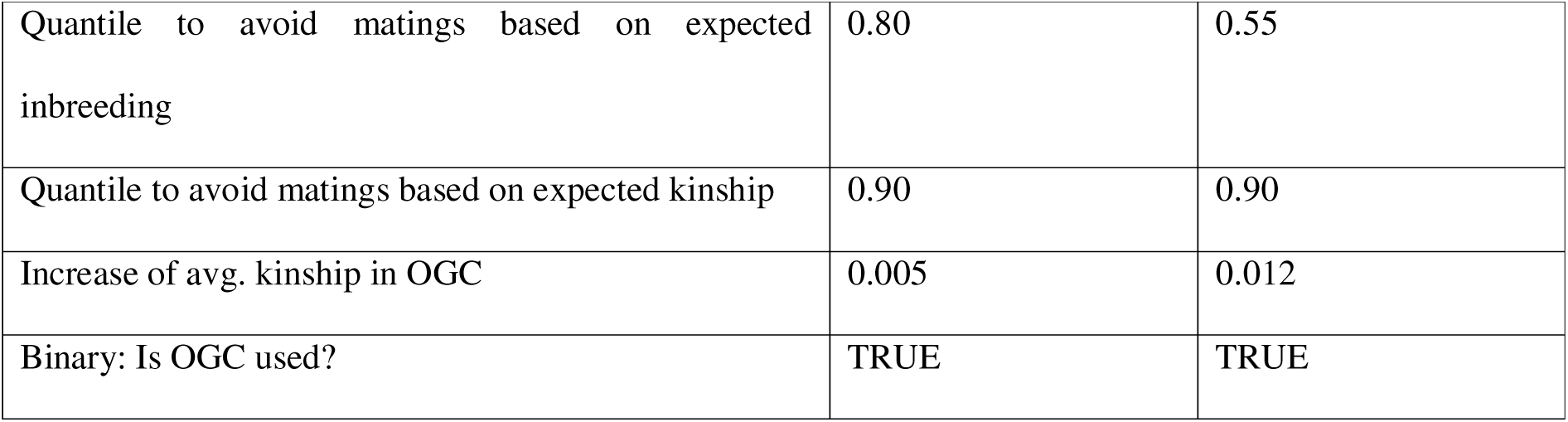
Suggested breeding program designs by the evolutionary pipeline [34] for the two potential target functions.

For both target functions, OGC is suggested to be used with a mild weighting of relatedness to top individuals in the selection index (17% / 9%). Furthermore, matings between related individuals are avoided, excluding 20% / 45% of all potential matings based on expected inbreeding, while only 10% of matings are avoided based on kinship in both scenarios. Furthermore, the evolutionary pipeline suggests strong emphasis on upweighting rare alleles by the moderate weighting of 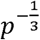, while limited weight is put on the theoretically optimal weight suggested by Goddard [20]. For both target functions, basically no index weight is put on the relatedness of an individual to the overall breeding population. The main difference between the two suggested breeding schemes is a much lower proportion of selected individuals in the scenario with focus on short-term gain, selecting 83 and 32 dams respectively (compared to 100 in the baseline).

The breeding program design optimized using the balanced target function leads to an increase in genetic gain of 1.34 gSD (+5.1%) and reduced inbreeding levels by 0.232 (-37.3%) after 50 generations. While gains relative to the baseline for inbreeding are obtained quite consistently over time, genetic gain for the first 20 generations is only on par with the reference scenario (Figures 10a and 10b).

**Figure 10:**
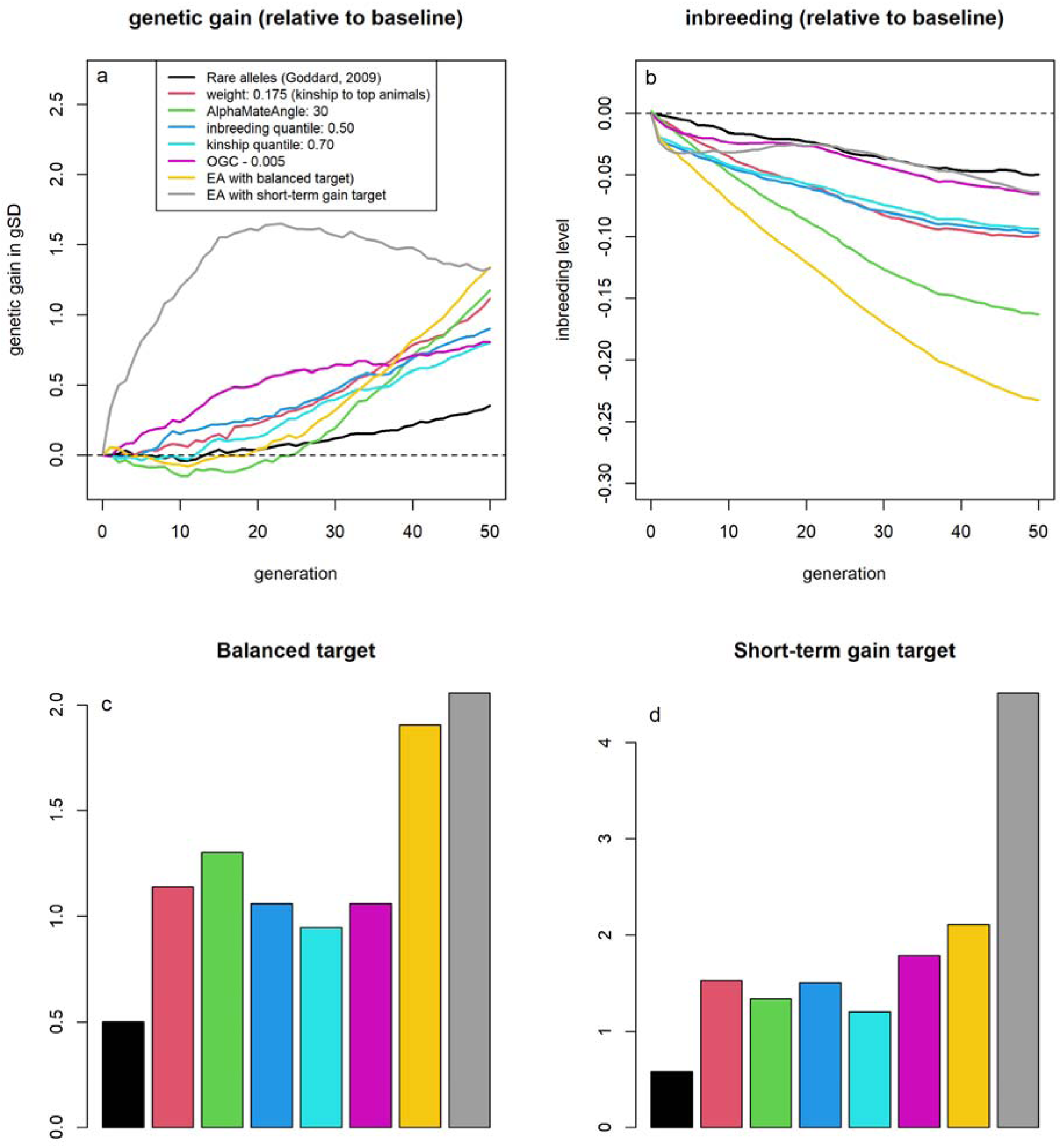
Genetic gain and inbreeding levels of scenarios optimized using the evolutionary algorithm. Genetic gain (a) and inbreeding levels (b) for various diversity management strategies relative to selection based on estimated breeding values. All scenarios are additionally evaluated on their performance on the balanced target function (c) and the target function with focus on short-term genetic gain (d) relative to selection based on estimated breeding values.

In contrast, the breeding program design optimized using the target function with focus on short-term genetic gain resulted in a reduction of inbreeding levels by 0.064 (-10.3%) and long-term genetic gains of 1.33 gSD (+5.1%). While reduction in inbreeding is obtained across the entire time horizon of the simulations, additional genetic gains are primary obtained in the short-term with an addition genetic gain of 0.33 gSD in the first generation (+31.1%; Figures 10a and 10b).

In comparison to breeding program designs that only make use of one of the management tools considered, the joint use of all techniques leads to much higher performance for the respective target. With the best individual technique resulting in an improvement of the target function of 1.30 / 1.15 relative to the baseline while the respective combination leading to an increase of the target function by 1.96 / 4.51 (Figures 10c and 10d).

## Discussion

This study provides an extensive comparison of potential selection strategies for the management of genetic diversity and inbreeding in a breeding program using stochastic simulations. In contrast to prior work, the focus of this study is less on developing new approaches to optimize selection in a breeding scheme, but on a comparison and assessment of existing techniques, and combinations thereof, within a unifying testing framework. Our results suggest the importance of careful assessment of individual approaches for management of genetic diversity in a breeding program to ensure their effectiveness. This is particularly true for breeding companies that have to find a balance between the maintenance of a healthy and diverse breeding population and the product they want to sell in the short-term [40].

### Individual selection strategies

The increased weighting of SNP effects of rare alleles with theoretically optimal long-term weights [20] yielded only net benefits in terms of genetic gain after 30 generations – which for dairy cattle would correspond to a time horizon of 75 years [41]. More moderate but still increased weighting of rare alleles (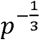) compared to Goddard [20] yielded marginally lower increases in long-term genetic gains while avoiding any short-term losses and therefore should be more practical for implementation in a commercial breeding program.

It is worth noting that the re-weighting of SNP effects based on allele frequencies overall only had a relatively low impact compared to other approaches tested. Although only a relatively small training population was considered for the prediction model, we do not expect major differences in larger populations as the use of real QTL effects for selection and allele weighting led to similar results with both higher short-term loss and long-term benefits (Supplementary Figure S7).

The use of the average kinship of individuals to the top individuals in the population as an additional trait in the selection index performed almost on par with the results obtained when using OGC [17]. For example, an index weight of 0.175 led to very similar genetic gain and inbreeding to use of OGC with a constraint of 0.43% inbreeding on the male side (Supplementary Figure S14), indicating very similar efficiency of approaches. Selection based on average kinship was even superior for long-term genetic gain. Furthermore, the use of a numeric score per individual provides substantial practical benefits compared to OGC as it is less computationally costly and a more tangible metric to work with which makes practical use very straightforward. On the other hand, selection based on average kinship lacks the ability to change short-term gain as possible in OGC by allowing for higher inbreeding rates.

In contrast to the previous studies [26, 28, 30], we did not find benefits in accounting for predicted progeny variance in the selection of individuals, that is potentially caused due to the breeding program design being unsuitable for the application of such strategies. First, selection proportion are high (top 8% males / 20% females) compared to the suggestions by Bijma et al. [25] that such strategies will only become effective with selection proportions of 5% or less. Second, no assortative mating strategies [26, 27, 30] were employed. Additionally, and perhaps most critically, the prediction accuracies for MSV in our study were low (0.17). Expanding the reference population improved prediction accuracies (0.30, Supplementary Figure 11), resulting in inbreeding and genetic gains comparable to those achieved through selecting based on EBVs.

Although not a technique traditionally associated with progeny variance, avoiding mating between related individuals should conceptually also lead to matings with a higher chance of new haplotypes and higher variation of potential offspring without the noise introduced by SNP effect estimation. The results here highlight very limited downsides of avoiding matings between individuals with high pairwise kinship and therefore high expected inbreeding levels of potential offspring.

From a practical breeding perspective many of the suggested approaches provide reduced inbreeding rates, but come at the cost of a reduced short-term genetic gain. However, these relative losses can be compensated by a decrease of selection proportions and/or use of a weaker inbreeding constraint in an OGC framework. Note here that an increase in genetic gain of 1% led to an increase in inbreeding rates by 4-5%. Therefore, a ratio between genetic gain and inbreeding as for example provided in [42, 43] should not be used as the sole metric for judging the long-term efficiency of a scenario as this will naturally favour approaches with lower genetic gain and inbreeding on a fixed time horizon. Instead one could for example consider changing the selection proportions to ensure similar inbreeding rates to make scenarios more comparable. Particularly for species with shorter generation interval like black soldier flies there is still value in the ratio as an additional metric of the outcomes of a breeding scheme [44] as in such schemes a state of basically no remaining genetic diversity and no further genetic progress can be reached, particularly in simulation.

#### Joint optimization

The suggested breeding scheme designs by the evolutionary algorithm [34] highlight that to maximize efficiency one should not limit oneself to a singular method for the management of genetic diversity, but use a combination of various approaches. The results of the evolutionary algorithm were naturally highly impacted by the actual choice of the target function. However, they still allow for some conclusions on which methods were the most efficient and which act complementary to each other. While weighting SNP effects by allele frequency of the beneficial allele had only limited overall effect, it acted complementary to OGC and the moderate weighting (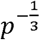) was chosen to be used in both runs of the evolutionary algorithm with substantial weighting. Although substantially lower selection proportions were used in both breeding program designs proposed by the evolutionary algorithm, the resulting inbreeding rates were actually lower than in the baseline scenario with selection based on estimated breeding values.

The evolutionary algorithm proposed by Hassanpour et al. [34] has been shown to be a powerful tool for the optimization of breeding programs, even though the initial search spaces did not contain the finally obtained optima as particularly our prior expectation on the weighting of rare alleles and selection intensity were off. From a practical perspective, we see two main challenges for the use of such methodology: Firstly, in the evolutionary algorithm 100 iterations with a total of 30,700 simulations each were performed, which for larger breeding schemes can require substantial computational costs and require the use of highly efficient simulation software [8, 10]. A potential solution could be to simplify the breeding program in a way that all considered scenarios should be affected in a similar way, e.g. to reduce computing time of a breeding value estimation by scaling individual numbers or reduce the number of traits considered. Note that, for computational reasons, the use of AlphaMate nor MSV were not considered in the optimization pipeline. Secondly, optimization requires a target to be optimized. Since assigning economic weights to traits can already be challenging in reality, we expect assigning economic weights of inbreeding rates and remaining genetic diversity to be even more difficult, while being of fundamental importance to initialize the optimization pipeline. The main solution we see for this is the use of experimental work with different target functions, and a subsequent analysis of the different optima obtained to derive their expected genetic gains and inbreeding and based on this decide which of the proposed optima is the most suitable in practice.

#### Practical consideration and implementation

Although the final optimum did contain a combination of various techniques, we fully acknowledge that from a practical perspective it can make sense to use a model that is as simple as possible (while getting most of the benefits) to avoid models becoming difficult to maintain and backpropagate. In this regard, it can of course make sense to reduce model complexity. In this regard, results on the individual strategies highlight which approaches to consider, with the use of MSV and weighting of SNP effects based on allele frequency being candidates to be dropped as their overall impact was low. Furthermore, the evolutionary optimization pipeline could be run with fewer parameters or the target function could include a penalty term for using more selection strategies.

As the breeding program considered in this study was chosen quite generic (with the strongest assumption being phenotyped and genotyped selection candidates), the reported impact of individual selection strategies should allow for general conclusions about their impact when used in a breeding program and allow for a pre-selection of methods to consider. For the fine-tuning of suitable weighting of each strategy, assessing stochastic simulations of a breeding program resembling the real-world breeding program (“digital twin”) can further improve the implementation.

Note here that the used trait was assumed to be purely additive and highly polygenic, without direct modelling on inbreeding depression [45], e.g., by including deleterious variants as dominant QTLs or epistasis [46, 47]. Hence, although inbreeding levels were increasing, genetic gain is only affected by limited genetic variance but not due to inbreeding depression. On the other hand, the genetic variance in stochastic simulations tends to decrease more than in reality – although in recent years this has also become more apparent in real-world breeding programs [48–50]. Potential reasons for this include that breeding programs in simulations are executed without error and also that in reality trait definition, breeding goals and environment might change over time [51–53]. Therefore, the results on the absolute levels of long-term genetic gain and genetic variance based on stochastic simulation should be taken with caution and conclusions should mainly be drawn based primarily on the relative ranking between scenarios. Here, one should critically question wheather different scenarios are affected in similar ways by the limitations and simplifications of the simulation.

This analysis by no means has claims of completeness and is mostly focused on the selection step itself and less on restructuring of the breeding program design as a whole to change generation interval [16], split the breeding scheme into multiple distinct components [15] or make use of cryo reserves [14] and outcross individuals [13]. All simulation and analysis scripts are provided in Supplementary File S15 and S16. Hereby, the flexibility of the simulation software MoBPS [8] allows for the extension of scripts to include further potential selection techniques and more general conceptional changes of breeding program design.

## Conclusions

The use of methods for the management of genetic diversity and inbreeding does not necessarily lead to short-term losses in term of genetic gain but can actually be used to simultaneously select less individuals.

The results in this study suggest that it is most efficient to use a combination of management strategies including OGC, weighting of SNP effects based on allele frequency, average kinship to top individuals as an additional trait in the selection index and avoiding mating between related individuals simultaneously and offset small short-term losses by selecting less individuals. As the use of all considered selection strategies comes at no additional costs other than additional computations, it is critical for breeding companies to implement such strategies for long-term success.

## Declarations

Ethics approval and consent to participate

## Not applicable

Consent for publication

## Not applicable

Availability of data and materials

The software MoBPS and the evolutionary pipeline can be found in the following two GitHub repositories: https://github.com/tpook92/MoBPS and https://github.com/AHassanpour88/Evolutionary_Snakemake. Data sharing is not applicable to this article as no datasets were generated or analysed during the current study.

## Authors’ contributions

TP led conceptualization, investigation and wrote the initial draft of the study. AH assisted on the adaptation of the evolutionary pipeline. TN and MC provided critical feedback in the investigation. MC acquired funding for the study. The first draft of the manuscript was written by TP. All authors reviewed, edited and approved the final manuscript.

## Acknowledgements

We acknowledge support, feedback, and discussions with Breed4Food partners in work package 4 (Genomic breeding program optimization).

## Additional files

Format: csv

Title: Supplementary File S1

Description: Genetic gain for all considered scenarios per generation.

Format: csv

Title: Supplementary File S2

Description: Inbreeding levels for all considered scenarios per generation.

Format: csv

Title: Supplementary File S3

Description: Remaining genetic variation compared to the first generation (based on genetic standard deviations of underlying true genomic values) for all considered scenario per generation.

Format: csv

Title: Supplementary File S4

Description: Share of SNPs fixated for all considered scenarios per generation.

Format: csv

Title: Supplementary File S5

Description: Share of QTLs fixated for all considered scenarios per generation.

Format: csv

Title: Supplementary File S6

Description: Share of QTLs fixated for the beneficial allele for all considered scenarios per generation.

**Supplementary Figure 7.**
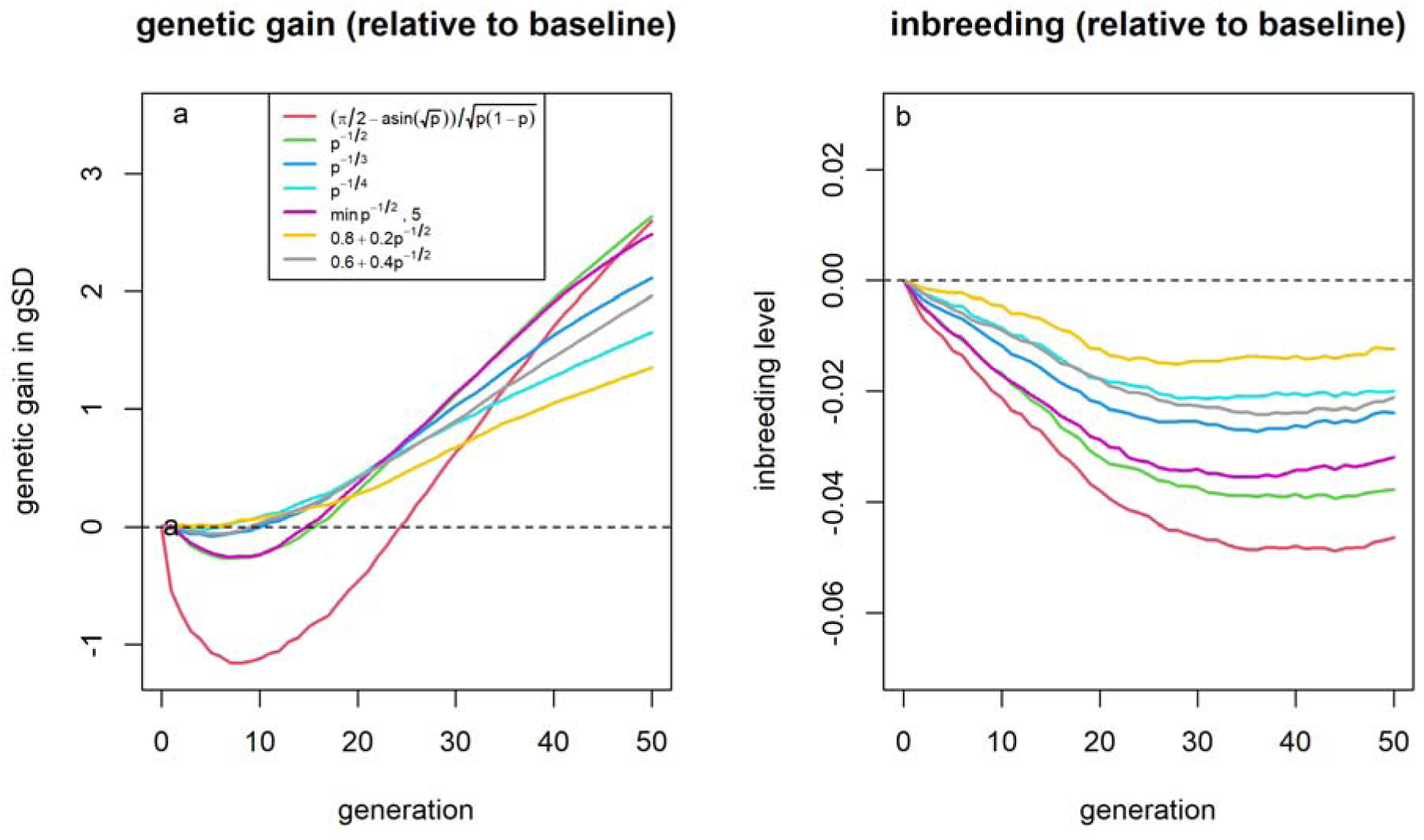
Genetic gains (a) and inbreeding levels (b) relative to selection based on underlying true genomic values for different weighting factors for SNP effects depending on the allele frequency of the beneficial variant, assuming known SNP effects.

**Supplementary Figure 8.**
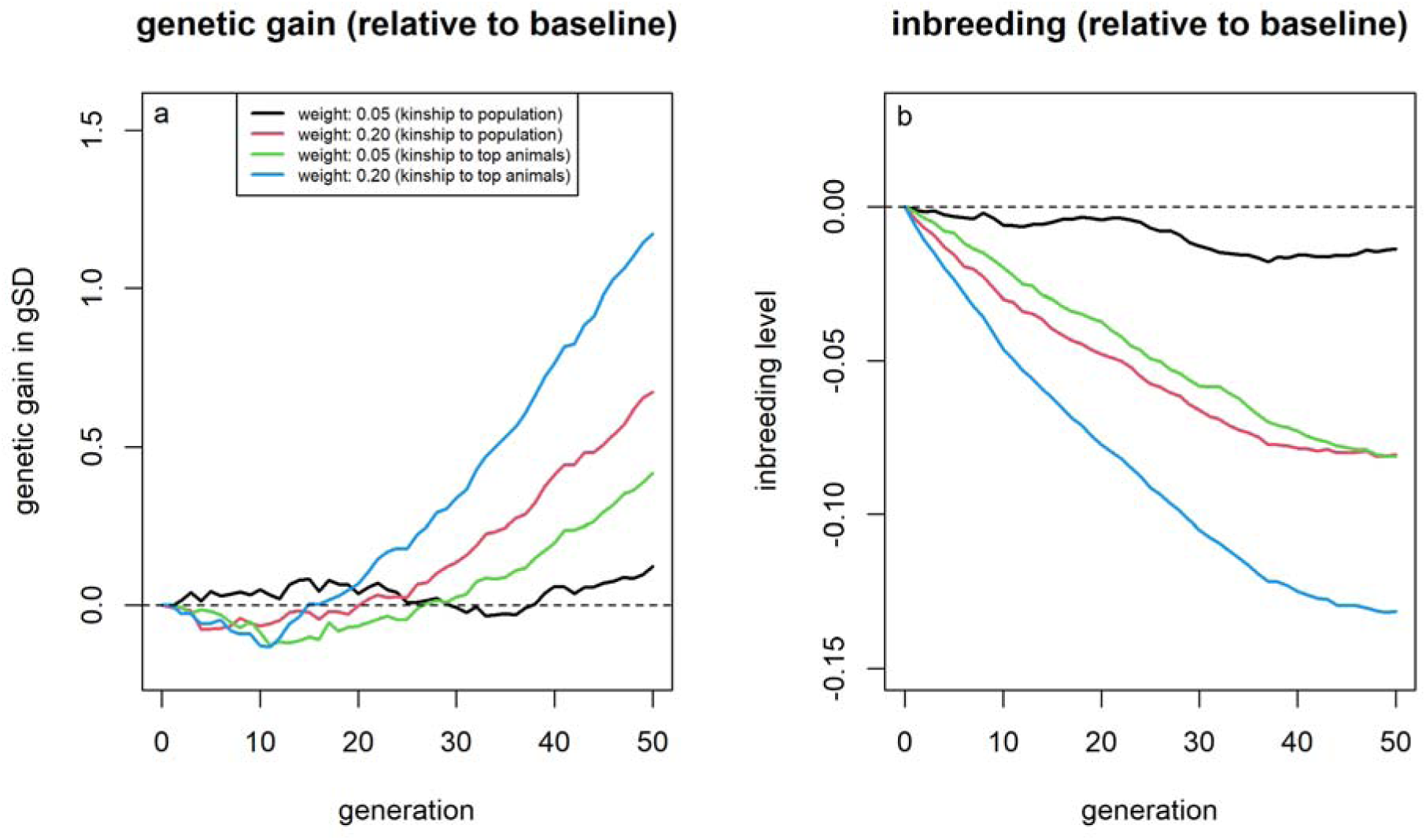
Genetic gains (a) and inbreeding levels (b) relative to selection based on estimated breeding values for different index weights for the average kinship of an individual to the current breeding population.

**Supplementary Figure 9.**
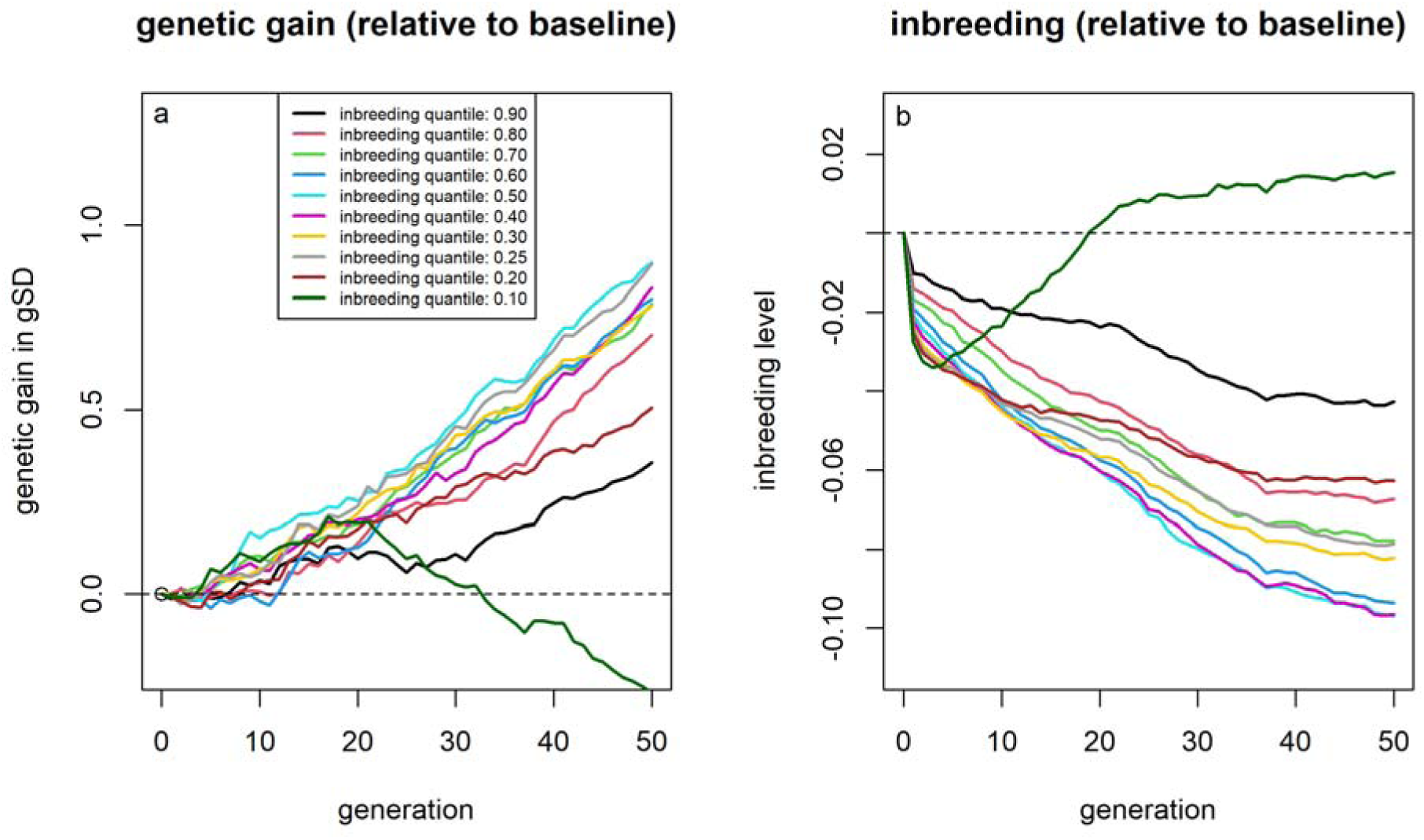
Genetic gains (a) and inbreeding levels (b) relative to selection based on estimated breeding values when avoiding different shares of matings between individuals based on the expected inbreeding level of a hypothetical offspring.

**Supplementary Figure 10.**
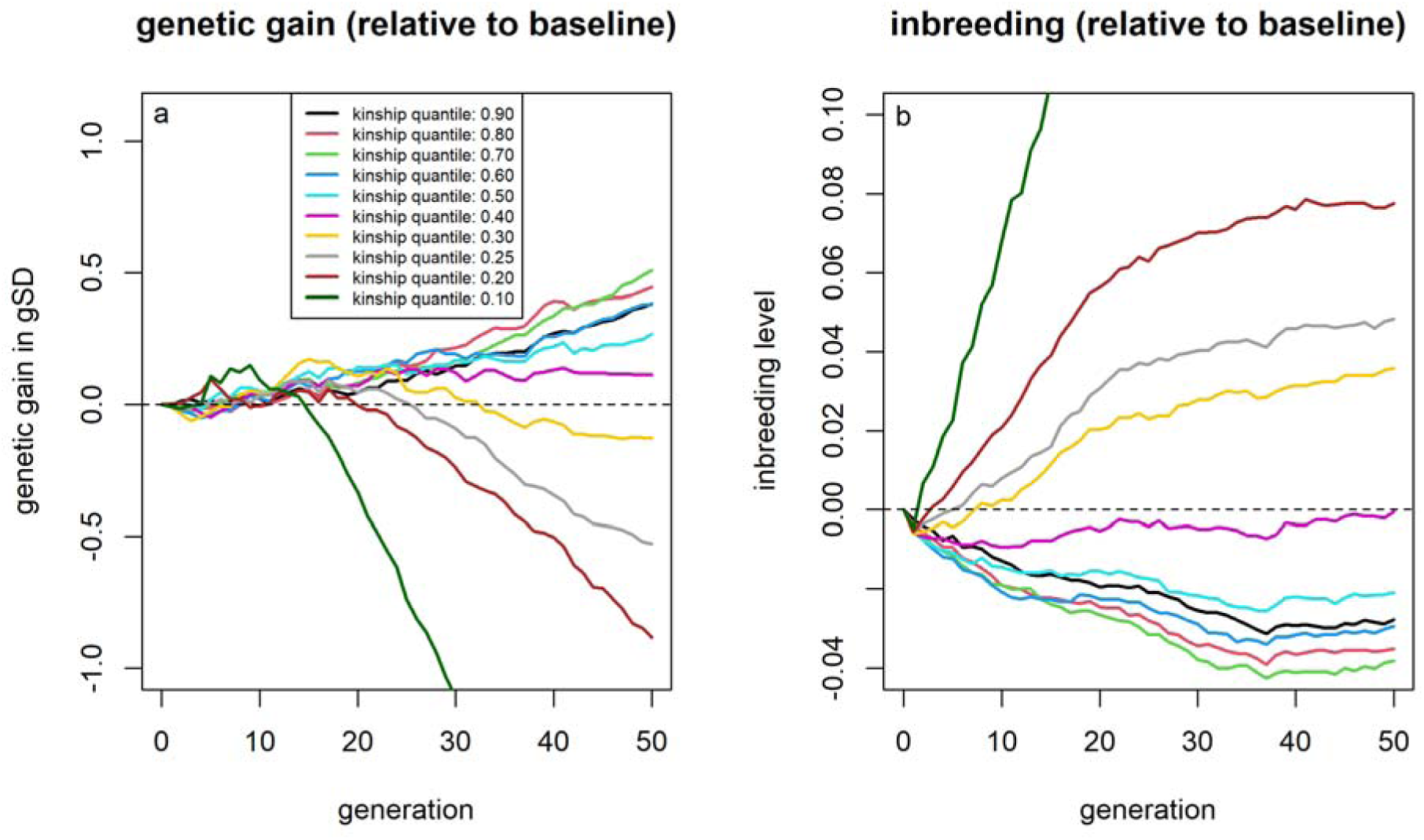
Genetic gains (a) and inbreeding levels (b) relative to selection based on estimated breeding values when avoiding different shares of matings between individuals based on the expected average kinship level to the current population of a hypothetical offspring.

**Supplementary Figure 11.**
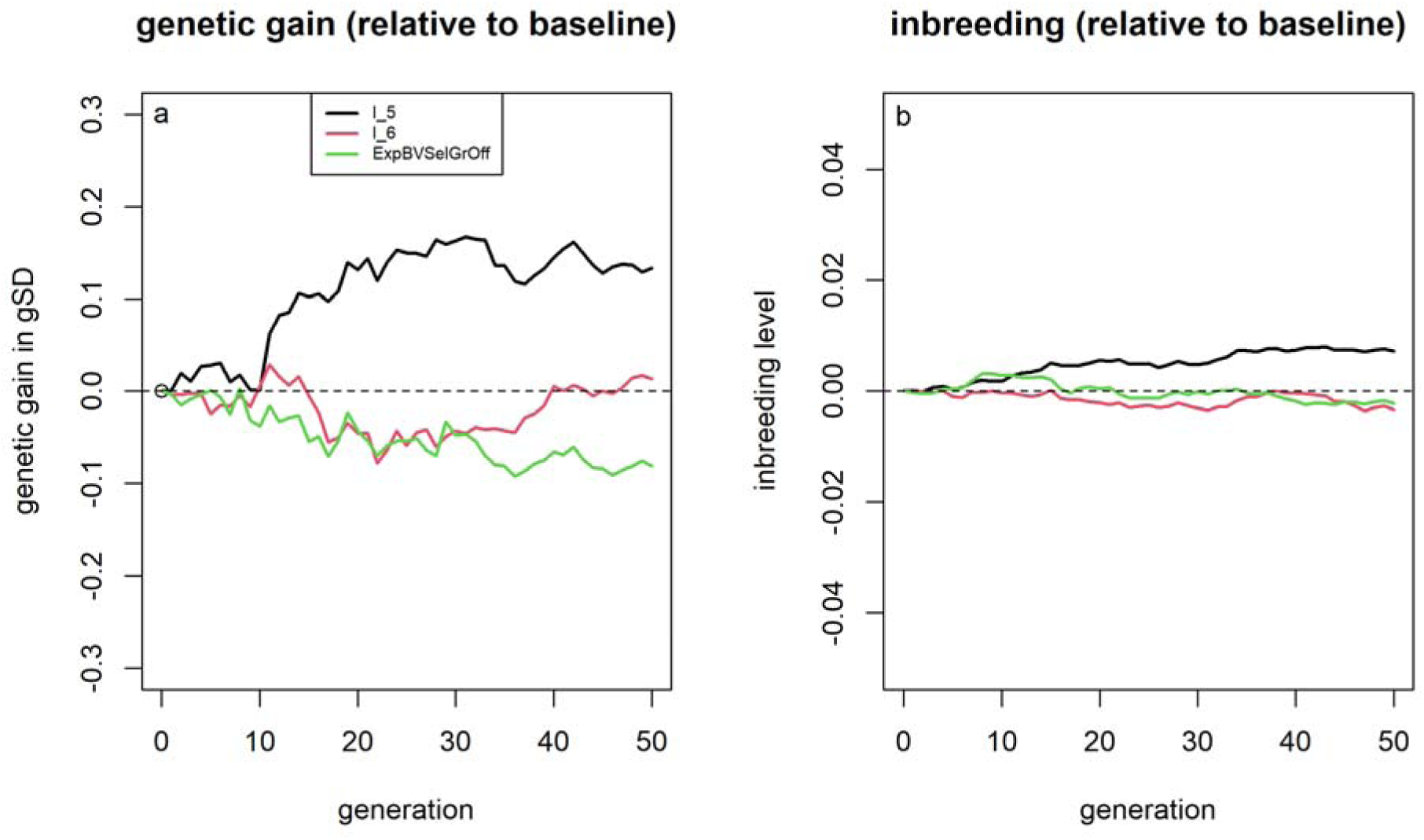
Genetic gains (a) and inbreeding levels (b) relative to selection based on estimated breeding values for different approaches to account for Mendelian sampling variance in selection when including the last six generations in the breeding value estimation.

**Supplementary Figure 12.**
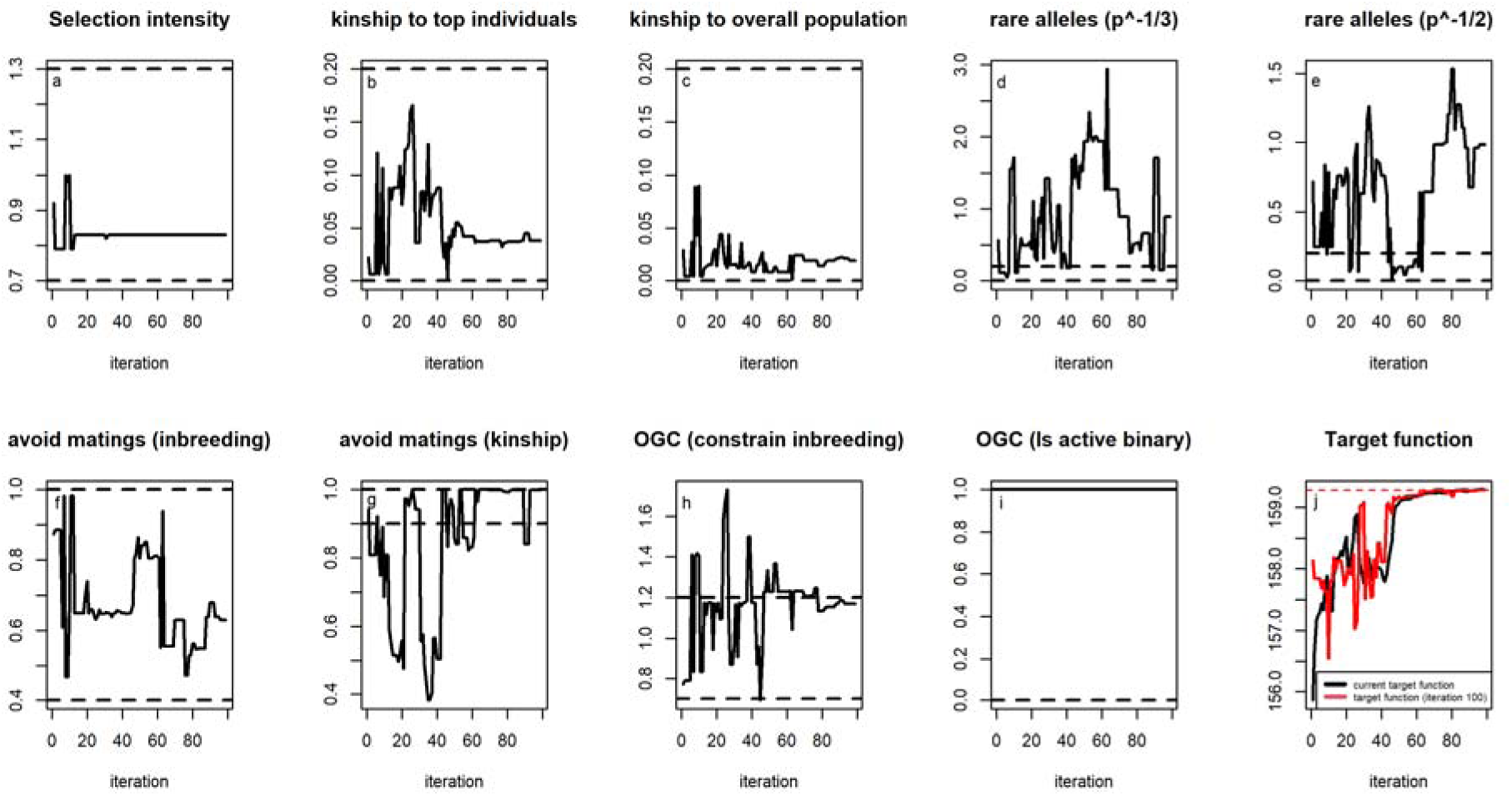
Optima suggested by the evolutionary algorithm with balanced target function for each iterations for each individual parameter and the estimated value of the target function in the optima. Black dashed lines indicate the initial sampling range per parameter.

**Supplementary Figure 13.**
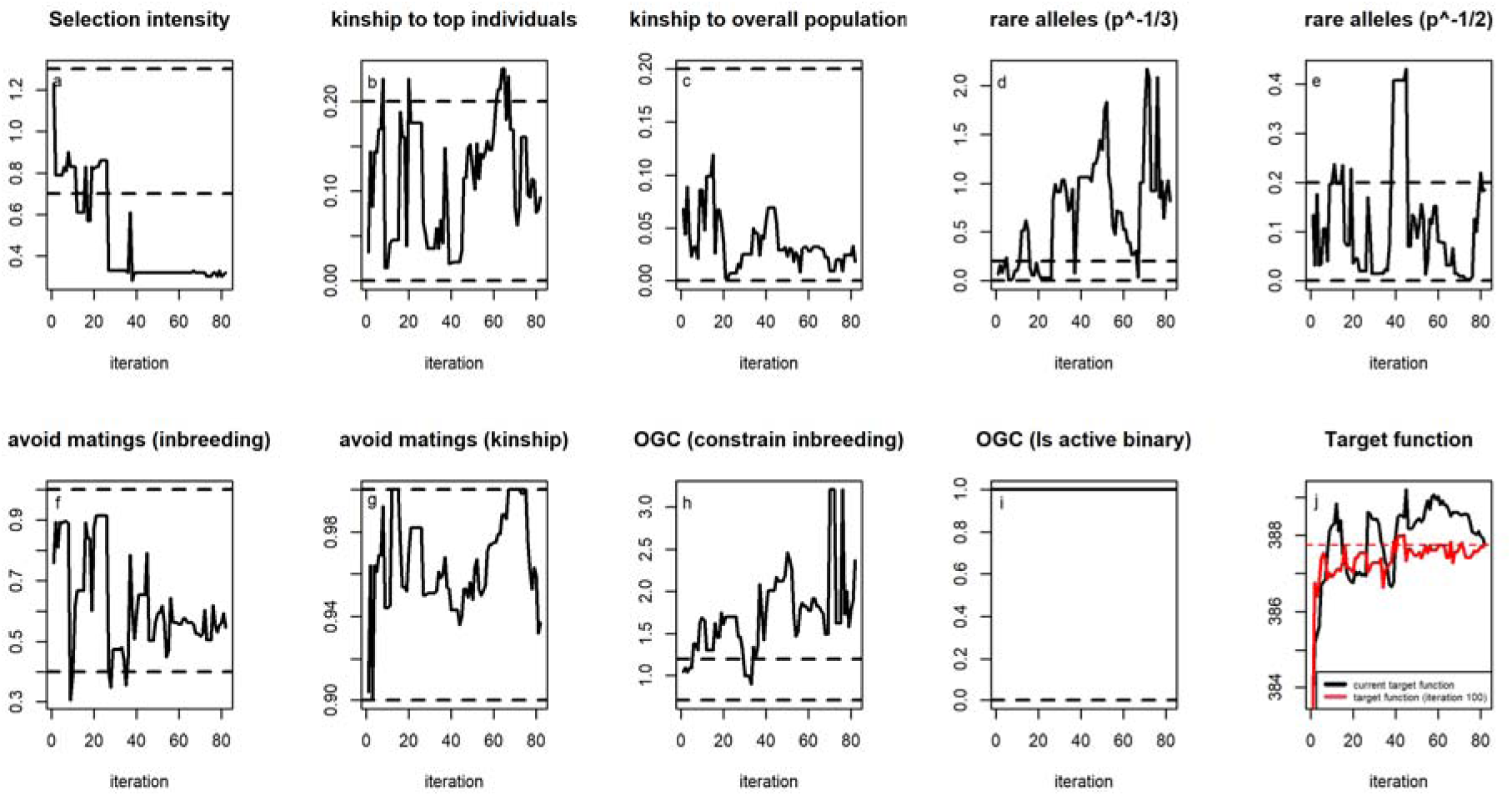
Optima suggested by the evolutionary algorithm with target function with focus on short-term genetic gain for each iterations for each individual parameter and the estimated value of the target function in the optima. Black dashed lines indicate the initial sampling range per parameter.

**Supplementary Figure 14.**
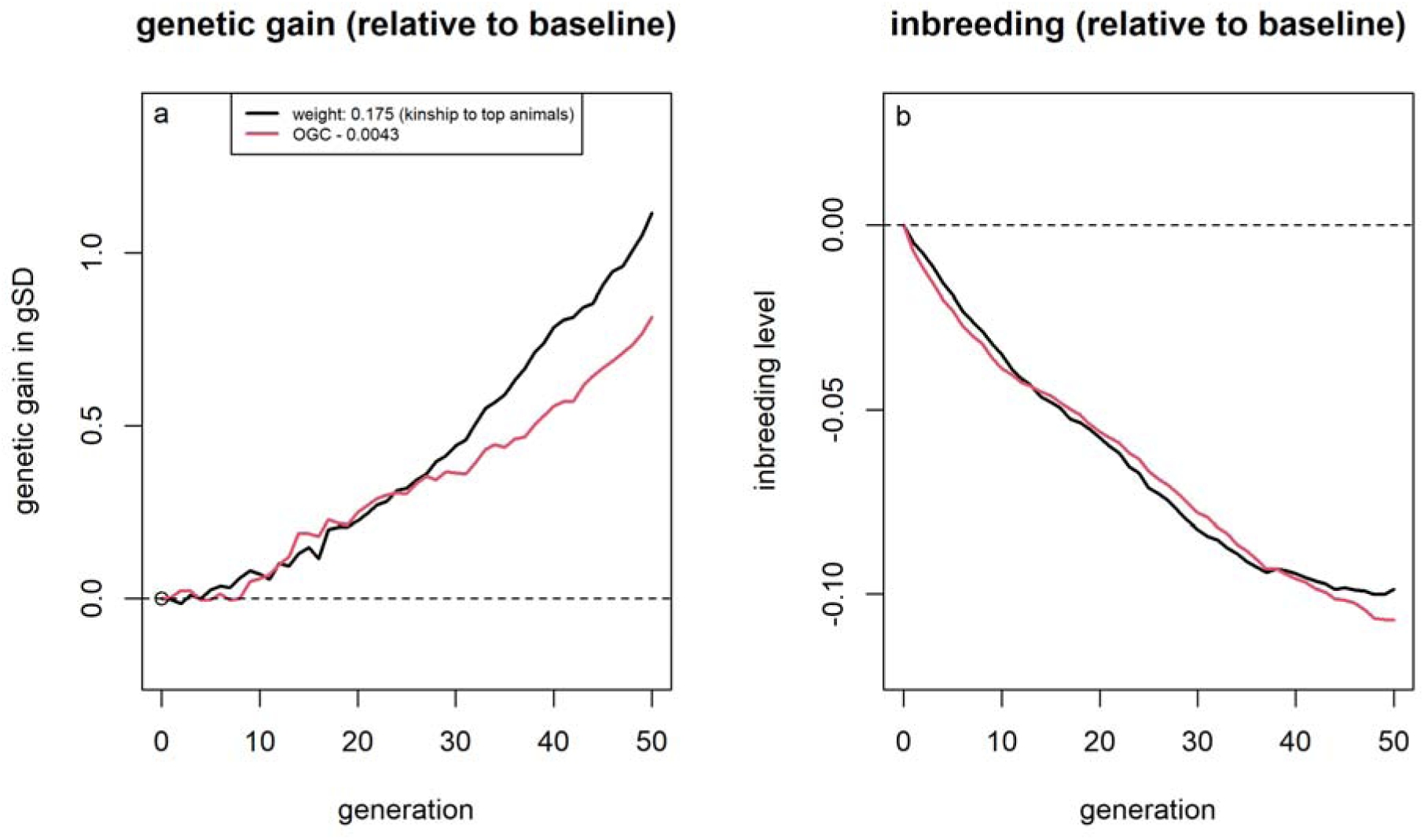
Genetic gains (a) and inbreeding levels (b) relative to selection based on estimated breeding values when using optimum genetic contribution selection with an maximum inbreeding of 0.43% compared to the use of average kinship to the top individuals as a trait in the selection index with a weight of 17.5%.

Format: R

Title: Supplementary File 15

Description: Script used for the simulation of the different breeding program design considered in this study and the subsequent analysis using the R-package MoBPS.

Format: R

Title: Supplementary File 16

Description: Script used for analysis of outcomes of the simulations and subsequent generation of plots.

