## Supplementary Files 15 &16 for "Strategies to improve selection compared to selection based on estimated breeding values": Supplementary_FileS15.R

```
#########################################################################################################
################################## Simulation script for manuscript: ####################################
#######  Strategies to improve selection  compared to selection based on estimated breeding values ######
#########################################################################################################

## Using args to submit different strategies within the same shell script
args <- commandArgs(TRUE)
scenario <- as.numeric(args[2])
seed <- as.numeric(args[1])

# some scenarios make use of currently WUR-internal software for estimation of
# Mendelian sampling variance. Formulas to derive MSV are given in:
# Niehoff et al. (https://gsejournal.biomedcentral.com/articles/10.1186/s12711-024-00899-2#additional-information)
# less efficient code is given in:
# Niehoff et al. (https://academic.oup.com/g3journal/article/14/11/jkae205/7743298?login=true#492554989)

# directory to write results to
title = paste0("uniqueness_v3/sc", scenario, "seed", seed, "_potential.RData")

library(MoBPS)

# Basic population parameters
n_additive = 1000 # Number of additive QTLs
heritability = 0.3 # heritability

pop_size = c(500, 500) # number of males / females per cycle
sel_size = c(40, 100) # number of selected males / females per cycle

nsnp = 25000 # number of SNPs
chr = 10 # number of chromosomes
chr_length = 2.5 # length of each chromosome in Morgan

burnin = 10
cycles = 50

weight_unique = 0.05

display.progress = FALSE

## Generation of the baseline population

set.seed(seed)

population = creating.diploid(nsnp = nsnp, nindi = sel_size,
                              chr.nr = chr, chromosome.length = chr_length,
                              n.additive = n_additive,
                              share.genotyped = 1)

alpha_real = numeric(nsnp)
alpha_real[population$info$real.bv.add[[1]][,6]] = (population$info$real.bv.add[[1]][,5] - population$info$real.bv.add[[1]][,3])/2


# Burn-in phase
population = breeding.diploid(population, heritability = heritability)
for(index in 1:burnin){
  population = breeding.diploid(population, phenotyping.gen = get.ngen(population))
  population = breeding.diploid(population, selection.size = sel_size, breeding.size = pop_size,
                                selection.criteria = "pheno",
                                share.genotyped = 1,
                                display.progress = display.progress)
}

# Standardization of traits in the first generation (after burn-in)
population = bv.standardization(population, mean.target = 100, var.target = 1,
                                gen = get.ngen(population),
                                adapt.pheno = TRUE)
population = breeding.diploid(population, heritability = heritability)

for(index in 1:cycles){
  
  current_gen = get.ngen(population)
  
  population = breeding.diploid(population, phenotyping.gen = current_gen)
  
  
  bve.gen = bve.gen1 = current_gen
  
  if(scenario %in% 335:338){
    bve.gen = (current_gen-5):current_gen
    bve.gen1  = current_gen
  }
  
  population = breeding.diploid(population, bve = TRUE, bve.gen = bve.gen, estimate.u = TRUE)
  
  alpha_hat = population$info$u_hat[[length(population$info$u_hat)]][,1] # estimated SNP effects
  
  if(sum(is.na(alpha_hat))>0){
    alpha_hat[is.na(alpha_hat)] = 0
  }
  geno = get.geno(population = population, gen = current_gen) # genotypes
  
  p = rowMeans(geno)/2
  
  # frequency of "good" allele
  freq_alpha = p
  freq_alpha[alpha_real==0] = 0 # these can be ignored
  freq_alpha[alpha_real<0] = 1 -freq_alpha[alpha_real<0] # these can be ignored
  
  # frequency of estimated "good" allele
  freq_alphahat = p
  freq_alphahat[alpha_hat==0] = 0 # these can be ignored
  freq_alphahat[alpha_hat<0] = 1 -freq_alphahat[alpha_hat<0] # these can be ignored
  
  ebv = get.bve(population, gen = current_gen)[1,] # estimated breeding value
  pheno = get.pheno(population, gen = current_gen)[1,] # phenotype
  bv = get.bv(population, gen = current_gen)[1,] # true underlying genomic value
  
  
  # this script was limited to those scenarios presented in the manuscript
  # scenario numbers were not changes (explaining why there is no scenario 2, 4, 6, 7, etc.)
  # order of scenarios corresponds to the other of results given in Supplementary Tables S1-S6
  
  if( scenario == 1){
    
    # Selection based on genomic breeding value
    population = breeding.diploid(population, selection.size = sel_size, breeding.size = pop_size,
                                  selection.criteria = "bve",
                                  share.genotyped = 1,
                                  display.progress = display.progress)
    
  } else if(scenario == 3){
    
    
    sel_size_tmp = sel_size
    # selection on the female side is done without OGC, on male male size selection.size = population size
    # to provide optiSel with all potential sires
    # calculation on how to set ub.sKin can be tricky has this corresponds to the increase of inbreeding
    # that is caused by the male side of the breeding program
    sel_size_tmp[1] = pop_size[1]
    # Selection based on genomic breeding value
    population = breeding.diploid(population, selection.size = sel_size_tmp,
                                  breeding.size = pop_size,
                                  selection.criteria = "bve",
                                  share.genotyped = 1,
                                  ogc = TRUE,
                                  ogc.target = "max.BV",
                                  ogc.uniform = "female",
                                  ogc.ub.sKin.increase = 0.005,
                                  display.progress = display.progress)
    
  } else if(scenario == 5){
    
    activ = alpha_hat!= 0
    
    weighting = sqrt(1 / freq_alphahat)
    weighting[weighting==Inf] = 0
    
    unique = as.numeric((alpha_hat[activ] * weighting[activ]) %*% geno[activ,])
    new_ebv = (unique - mean(unique))/sd(unique)
    
    population = insert.bve(population, bves = cbind(names(ebv), new_ebv))
    population = breeding.diploid(population, selection.size = sel_size, breeding.size = pop_size,
                                  selection.criteria = "bve",
                                  share.genotyped = 1,
                                  display.progress = display.progress)
    
  } else if( scenario == 10){
    
    
    cand = breeding.diploid(population, selection.size = sel_size, breeding.size = pop_size,
                            selection.criteria = "bve", export.selected = TRUE )
    
    A = kinship.exp(population, gen = get.ngen(population))
    
    rel_top = colMeans(A[cand[[1]][,3],]) + colMeans(A[cand[[2]][,3] + pop_size[1],])
    
    unique = -rel_top #
    new_ebv = weight_unique * (unique - mean(unique))/sd(unique) + (1-weight_unique)* (ebv - mean(ebv))/sd(ebv)
    
    population = insert.bve(population, bves = cbind(names(ebv), new_ebv))
    population = breeding.diploid(population, selection.size = sel_size, breeding.size = pop_size,
                                  selection.criteria = "bve",
                                  share.genotyped = 1,
                                  display.progress = display.progress)
  } else if( scenario == 14){
    
    weight_unique = 0.2
    
    cand = breeding.diploid(population, selection.size = sel_size, breeding.size = pop_size,
                            selection.criteria = "bve", export.selected = TRUE )
    
    A = kinship.exp(population, gen = get.ngen(population))
    
    rel_top = colMeans(A[cand[[1]][,3],]) + colMeans(A[cand[[2]][,3] + pop_size[1],])
    
    unique = -rel_top #
    new_ebv = weight_unique * (unique - mean(unique))/sd(unique) + (1-weight_unique)* (ebv - mean(ebv))/sd(ebv)
    
    population = insert.bve(population, bves = cbind(names(ebv), new_ebv))
    population = breeding.diploid(population, selection.size = sel_size, breeding.size = pop_size,
                                  selection.criteria = "bve",
                                  share.genotyped = 1,
                                  display.progress = display.progress)
  } else if( scenario == 36){
    
    cand = breeding.diploid(population, selection.size = sel_size, breeding.size = pop_size,
                            selection.criteria = "bve", export.selected = TRUE )
    
    A = kinship.exp(population, gen = get.ngen(population))
    
    rel_top = colMeans(A)
    
    unique = -rel_top #
    new_ebv = weight_unique * (unique - mean(unique))/sd(unique) + (1-weight_unique)* (ebv - mean(ebv))/sd(ebv)
    
    population = insert.bve(population, bves = cbind(names(ebv), new_ebv))
    population = breeding.diploid(population, selection.size = sel_size, breeding.size = pop_size,
                                  selection.criteria = "bve",
                                  share.genotyped = 1,
                                  display.progress = display.progress)
  } else if( scenario == 37){
    
    weight_unique = 0.2
    
    cand = breeding.diploid(population, selection.size = sel_size, breeding.size = pop_size,
                            selection.criteria = "bve", export.selected = TRUE )
    
    A = kinship.exp(population, gen = get.ngen(population))
    
    rel_top = colMeans(A)
    
    unique = -rel_top #
    new_ebv = weight_unique * (unique - mean(unique))/sd(unique) + (1-weight_unique)* (ebv - mean(ebv))/sd(ebv)
    
    population = insert.bve(population, bves = cbind(names(ebv), new_ebv))
    population = breeding.diploid(population, selection.size = sel_size, breeding.size = pop_size,
                                  selection.criteria = "bve",
                                  share.genotyped = 1,
                                  display.progress = display.progress)
  } else if( scenario == 53){
    
    weight_unique = 0.025
    cand = breeding.diploid(population, selection.size = sel_size, breeding.size = pop_size,
                            selection.criteria = "bve", export.selected = TRUE )
    
    A = kinship.exp(population, gen = get.ngen(population))
    diag(A) = (diag(A)-0.5)*2
    
    rel_top = colMeans(A[cand[[1]][,3],]) + colMeans(A[cand[[2]][,3] + pop_size[1],])
    
    unique = -rel_top #
    new_ebv = weight_unique * (unique - mean(unique))/sd(unique) + (1-weight_unique)* (ebv - mean(ebv))/sd(ebv)
    
    population = insert.bve(population, bves = cbind(names(ebv), new_ebv))
    population = breeding.diploid(population, selection.size = sel_size, breeding.size = pop_size,
                                  selection.criteria = "bve",
                                  share.genotyped = 1,
                                  display.progress = display.progress)
  } else if( scenario == 54){
    
    weight_unique = 0.075
    cand = breeding.diploid(population, selection.size = sel_size, breeding.size = pop_size,
                            selection.criteria = "bve", export.selected = TRUE )
    
    A = kinship.exp(population, gen = get.ngen(population))
    diag(A) = (diag(A)-0.5)*2
    
    rel_top = colMeans(A[cand[[1]][,3],]) + colMeans(A[cand[[2]][,3] + pop_size[1],])
    
    unique = -rel_top #
    new_ebv = weight_unique * (unique - mean(unique))/sd(unique) + (1-weight_unique)* (ebv - mean(ebv))/sd(ebv)
    
    population = insert.bve(population, bves = cbind(names(ebv), new_ebv))
    population = breeding.diploid(population, selection.size = sel_size, breeding.size = pop_size,
                                  selection.criteria = "bve",
                                  share.genotyped = 1,
                                  display.progress = display.progress)
  } else if( scenario == 55){
    
    weight_unique = 0.1
    cand = breeding.diploid(population, selection.size = sel_size, breeding.size = pop_size,
                            selection.criteria = "bve", export.selected = TRUE )
    
    A = kinship.exp(population, gen = get.ngen(population))
    diag(A) = (diag(A)-0.5)*2
    
    rel_top = colMeans(A[cand[[1]][,3],]) + colMeans(A[cand[[2]][,3] + pop_size[1],])
    
    unique = -rel_top #
    new_ebv = weight_unique * (unique - mean(unique))/sd(unique) + (1-weight_unique)* (ebv - mean(ebv))/sd(ebv)
    
    population = insert.bve(population, bves = cbind(names(ebv), new_ebv))
    population = breeding.diploid(population, selection.size = sel_size, breeding.size = pop_size,
                                  selection.criteria = "bve",
                                  share.genotyped = 1,
                                  display.progress = display.progress)
  } else if( scenario == 56){
    
    weight_unique = 0.125
    cand = breeding.diploid(population, selection.size = sel_size, breeding.size = pop_size,
                            selection.criteria = "bve", export.selected = TRUE )
    
    A = kinship.exp(population, gen = get.ngen(population))
    diag(A) = (diag(A)-0.5)*2
    
    rel_top = colMeans(A[cand[[1]][,3],]) + colMeans(A[cand[[2]][,3] + pop_size[1],])
    
    unique = -rel_top #
    new_ebv = weight_unique * (unique - mean(unique))/sd(unique) + (1-weight_unique)* (ebv - mean(ebv))/sd(ebv)
    
    population = insert.bve(population, bves = cbind(names(ebv), new_ebv))
    population = breeding.diploid(population, selection.size = sel_size, breeding.size = pop_size,
                                  selection.criteria = "bve",
                                  share.genotyped = 1,
                                  display.progress = display.progress)
  }else if( scenario == 57){
    
    weight_unique = 0.15
    cand = breeding.diploid(population, selection.size = sel_size, breeding.size = pop_size,
                            selection.criteria = "bve", export.selected = TRUE )
    
    A = kinship.exp(population, gen = get.ngen(population))
    diag(A) = (diag(A)-0.5)*2
    
    rel_top = colMeans(A[cand[[1]][,3],]) + colMeans(A[cand[[2]][,3] + pop_size[1],])
    
    unique = -rel_top #
    new_ebv = weight_unique * (unique - mean(unique))/sd(unique) + (1-weight_unique)* (ebv - mean(ebv))/sd(ebv)
    
    population = insert.bve(population, bves = cbind(names(ebv), new_ebv))
    population = breeding.diploid(population, selection.size = sel_size, breeding.size = pop_size,
                                  selection.criteria = "bve",
                                  share.genotyped = 1,
                                  display.progress = display.progress)
  } else if( scenario == 58){
    
    weight_unique = 0.175
    cand = breeding.diploid(population, selection.size = sel_size, breeding.size = pop_size,
                            selection.criteria = "bve", export.selected = TRUE )
    
    A = kinship.exp(population, gen = get.ngen(population))
    diag(A) = (diag(A)-0.5)*2
    
    rel_top = colMeans(A[cand[[1]][,3],]) + colMeans(A[cand[[2]][,3] + pop_size[1],])
    
    unique = -rel_top #
    new_ebv = weight_unique * (unique - mean(unique))/sd(unique) + (1-weight_unique)* (ebv - mean(ebv))/sd(ebv)
    
    population = insert.bve(population, bves = cbind(names(ebv), new_ebv))
    population = breeding.diploid(population, selection.size = sel_size, breeding.size = pop_size,
                                  selection.criteria = "bve",
                                  share.genotyped = 1,
                                  display.progress = display.progress)
  } else if( scenario == 59){
    
    weight_unique = 0.25
    cand = breeding.diploid(population, selection.size = sel_size, breeding.size = pop_size,
                            selection.criteria = "bve", export.selected = TRUE )
    
    A = kinship.exp(population, gen = get.ngen(population))
    diag(A) = (diag(A)-0.5)*2
    
    rel_top = colMeans(A[cand[[1]][,3],]) + colMeans(A[cand[[2]][,3] + pop_size[1],])
    
    unique = -rel_top #
    new_ebv = weight_unique * (unique - mean(unique))/sd(unique) + (1-weight_unique)* (ebv - mean(ebv))/sd(ebv)
    
    population = insert.bve(population, bves = cbind(names(ebv), new_ebv))
    population = breeding.diploid(population, selection.size = sel_size, breeding.size = pop_size,
                                  selection.criteria = "bve",
                                  share.genotyped = 1,
                                  display.progress = display.progress)
  }else if( scenario == 60){
    
    weight_unique = 0.3
    cand = breeding.diploid(population, selection.size = sel_size, breeding.size = pop_size,
                            selection.criteria = "bve", export.selected = TRUE )
    
    A = kinship.exp(population, gen = get.ngen(population))
    diag(A) = (diag(A)-0.5)*2
    
    rel_top = colMeans(A[cand[[1]][,3],]) + colMeans(A[cand[[2]][,3] + pop_size[1],])
    
    unique = -rel_top #
    new_ebv = weight_unique * (unique - mean(unique))/sd(unique) + (1-weight_unique)* (ebv - mean(ebv))/sd(ebv)
    
    population = insert.bve(population, bves = cbind(names(ebv), new_ebv))
    population = breeding.diploid(population, selection.size = sel_size, breeding.size = pop_size,
                                  selection.criteria = "bve",
                                  share.genotyped = 1,
                                  display.progress = display.progress)
  } else if( scenario == 61){
    
    weight_unique = 0.4
    cand = breeding.diploid(population, selection.size = sel_size, breeding.size = pop_size,
                            selection.criteria = "bve", export.selected = TRUE )
    
    A = kinship.exp(population, gen = get.ngen(population))
    diag(A) = (diag(A)-0.5)*2
    
    rel_top = colMeans(A[cand[[1]][,3],]) + colMeans(A[cand[[2]][,3] + pop_size[1],])
    
    unique = -rel_top #
    new_ebv = weight_unique * (unique - mean(unique))/sd(unique) + (1-weight_unique)* (ebv - mean(ebv))/sd(ebv)
    
    population = insert.bve(population, bves = cbind(names(ebv), new_ebv))
    population = breeding.diploid(population, selection.size = sel_size, breeding.size = pop_size,
                                  selection.criteria = "bve",
                                  share.genotyped = 1,
                                  display.progress = display.progress)
  } else if( scenario == 62){
    
    weight_unique = 0.5
    cand = breeding.diploid(population, selection.size = sel_size, breeding.size = pop_size,
                            selection.criteria = "bve", export.selected = TRUE )
    
    A = kinship.exp(population, gen = get.ngen(population))
    diag(A) = (diag(A)-0.5)*2
    
    rel_top = colMeans(A[cand[[1]][,3],]) + colMeans(A[cand[[2]][,3] + pop_size[1],])
    
    unique = -rel_top #
    new_ebv = weight_unique * (unique - mean(unique))/sd(unique) + (1-weight_unique)* (ebv - mean(ebv))/sd(ebv)
    
    population = insert.bve(population, bves = cbind(names(ebv), new_ebv))
    population = breeding.diploid(population, selection.size = sel_size, breeding.size = pop_size,
                                  selection.criteria = "bve",
                                  share.genotyped = 1,
                                  display.progress = display.progress)
  } else if( scenario == 95){
    
    # Selection based on genomic breeding value
    population = breeding.diploid(population, selection.size = sel_size, breeding.size = pop_size,
                                  selection.criteria = "bve",
                                  share.genotyped = 1,
                                  avoid.mating.fullsib = TRUE,
                                  display.progress = display.progress)
    
  } else if( scenario == 96){
    
    # Selection based on genomic breeding value
    population = breeding.diploid(population, selection.size = sel_size, breeding.size = pop_size,
                                  selection.criteria = "bve",
                                  share.genotyped = 1,
                                  avoid.mating.halfsib = TRUE,
                                  display.progress = display.progress)
    
  } else if( scenario == 97){
    
    # Selection based on genomic breeding value
    population = breeding.diploid(population, selection.size = sel_size, breeding.size = pop_size,
                                  selection.criteria = "bve",
                                  share.genotyped = 1,
                                  avoid.mating.inb.quantile = 0.9,
                                  display.progress = display.progress)
    
  } else if( scenario == 98){
    
    # Selection based on genomic breeding value
    population = breeding.diploid(population, selection.size = sel_size, breeding.size = pop_size,
                                  selection.criteria = "bve",
                                  share.genotyped = 1,
                                  avoid.mating.inb.quantile = 0.5,
                                  display.progress = display.progress)
    
  } else if( scenario == 99){
    
    # Selection based on genomic breeding value
    population = breeding.diploid(population, selection.size = sel_size, breeding.size = pop_size,
                                  selection.criteria = "bve",
                                  share.genotyped = 1,
                                  avoid.mating.inb.quantile = 0.25,
                                  display.progress = display.progress)
    
  } else if( scenario == 100){
    
    # Selection based on genomic breeding value
    population = breeding.diploid(population, selection.size = sel_size, breeding.size = pop_size,
                                  selection.criteria = "bve",
                                  share.genotyped = 1,
                                  avoid.mating.inb.quantile = 0.1,
                                  display.progress = display.progress)
    
  }  else if( scenario == 118){
    
    # Selection based on genomic breeding value
    population = breeding.diploid(population, selection.size = sel_size, breeding.size = pop_size,
                                  selection.criteria = "bve",
                                  share.genotyped = 1,
                                  avoid.mating.kinship.quantile = 0.9,
                                  avoid.mating.kinship.gen = get.ngen(population),
                                  display.progress = display.progress)
    
  } else if( scenario == 119){
    
    # Selection based on genomic breeding value
    population = breeding.diploid(population, selection.size = sel_size, breeding.size = pop_size,
                                  selection.criteria = "bve",
                                  share.genotyped = 1,
                                  avoid.mating.kinship.quantile = 0.5,
                                  avoid.mating.kinship.gen = get.ngen(population),
                                  display.progress = display.progress)
    
  } else if( scenario == 120){
    
    # Selection based on genomic breeding value
    population = breeding.diploid(population, selection.size = sel_size, breeding.size = pop_size,
                                  selection.criteria = "bve",
                                  share.genotyped = 1,
                                  avoid.mating.kinship.quantile = 0.25,
                                  avoid.mating.kinship.gen = get.ngen(population),
                                  display.progress = display.progress)
    
  } else if( scenario == 121){
    
    # Selection based on genomic breeding value
    population = breeding.diploid(population, selection.size = sel_size, breeding.size = pop_size,
                                  selection.criteria = "bve",
                                  share.genotyped = 1,
                                  avoid.mating.kinship.quantile = 0.1,
                                  avoid.mating.kinship.gen = get.ngen(population),
                                  display.progress = display.progress)
    
  } else if( scenario == 125){
    
    # Selection based on genomic breeding value
    population = breeding.diploid(population, selection.size = sel_size, breeding.size = pop_size,
                                  selection.criteria = "bve",
                                  share.genotyped = 1,
                                  avoid.mating.inb.quantile = 0.8,
                                  display.progress = display.progress)
    
  } else if( scenario == 126){
    
    # Selection based on genomic breeding value
    population = breeding.diploid(population, selection.size = sel_size, breeding.size = pop_size,
                                  selection.criteria = "bve",
                                  share.genotyped = 1,
                                  avoid.mating.inb.quantile = 0.7,
                                  display.progress = display.progress)
    
  } else if( scenario == 127){
    
    # Selection based on genomic breeding value
    population = breeding.diploid(population, selection.size = sel_size, breeding.size = pop_size,
                                  selection.criteria = "bve",
                                  share.genotyped = 1,
                                  avoid.mating.inb.quantile = 0.6,
                                  display.progress = display.progress)
    
  } else if( scenario == 130){
    
    # Selection based on genomic breeding value
    population = breeding.diploid(population, selection.size = sel_size, breeding.size = pop_size,
                                  selection.criteria = "bve",
                                  share.genotyped = 1,
                                  avoid.mating.inb.quantile = 0.4,
                                  display.progress = display.progress)
    
  } else if( scenario == 131){
    
    # Selection based on genomic breeding value
    population = breeding.diploid(population, selection.size = sel_size, breeding.size = pop_size,
                                  selection.criteria = "bve",
                                  share.genotyped = 1,
                                  avoid.mating.inb.quantile = 0.3,
                                  display.progress = display.progress)
    
  } else if( scenario == 132){
    
    # Selection based on genomic breeding value
    population = breeding.diploid(population, selection.size = sel_size, breeding.size = pop_size,
                                  selection.criteria = "bve",
                                  share.genotyped = 1,
                                  avoid.mating.inb.quantile = 0.2,
                                  display.progress = display.progress)
    
  } else if( scenario == 133){
    
    # Selection based on genomic breeding value
    population = breeding.diploid(population, selection.size = sel_size, breeding.size = pop_size,
                                  selection.criteria = "bve",
                                  share.genotyped = 1,
                                  avoid.mating.kinship.quantile = 0.8,
                                  avoid.mating.kinship.gen = get.ngen(population),
                                  display.progress = display.progress)
    
  } else if( scenario == 134){
    
    # Selection based on genomic breeding value
    population = breeding.diploid(population, selection.size = sel_size, breeding.size = pop_size,
                                  selection.criteria = "bve",
                                  share.genotyped = 1,
                                  avoid.mating.kinship.quantile = 0.7,
                                  avoid.mating.kinship.gen = get.ngen(population),
                                  display.progress = display.progress)
    
  } else if( scenario == 135){
    
    # Selection based on genomic breeding value
    population = breeding.diploid(population, selection.size = sel_size, breeding.size = pop_size,
                                  selection.criteria = "bve",
                                  share.genotyped = 1,
                                  avoid.mating.kinship.quantile = 0.6,
                                  avoid.mating.kinship.gen = get.ngen(population),
                                  display.progress = display.progress)
    
  }else if( scenario == 136){
    
    # Selection based on genomic breeding value
    population = breeding.diploid(population, selection.size = sel_size, breeding.size = pop_size,
                                  selection.criteria = "bve",
                                  share.genotyped = 1,
                                  avoid.mating.kinship.quantile = 0.4,
                                  avoid.mating.kinship.gen = get.ngen(population),
                                  display.progress = display.progress)
    
  }else if( scenario == 137){
    
    # Selection based on genomic breeding value
    population = breeding.diploid(population, selection.size = sel_size, breeding.size = pop_size,
                                  selection.criteria = "bve",
                                  share.genotyped = 1,
                                  avoid.mating.kinship.quantile = 0.3,
                                  avoid.mating.kinship.gen = get.ngen(population),
                                  display.progress = display.progress)
    
  }else if( scenario == 138){
    
    # Selection based on genomic breeding value
    population = breeding.diploid(population, selection.size = sel_size, breeding.size = pop_size,
                                  selection.criteria = "bve",
                                  share.genotyped = 1,
                                  avoid.mating.kinship.quantile = 0.2,
                                  avoid.mating.kinship.gen = get.ngen(population),
                                  display.progress = display.progress)
    
  } else if(scenario == 187){
    
    activ = alpha_hat!= 0
    
    weighting = (1 / freq_alphahat)^(1/2)
    weighting[weighting==Inf] = 0
    
    unique = as.numeric((alpha_hat[activ] * weighting[activ]) %*% geno[activ,])
    new_ebv = (unique - mean(unique))/sd(unique)
    
    population = insert.bve(population, bves = cbind(names(ebv), new_ebv))
    population = breeding.diploid(population, selection.size = sel_size, breeding.size = pop_size,
                                  selection.criteria = "bve",
                                  share.genotyped = 1,
                                  display.progress = display.progress)
    
  } else if(scenario == 188){
    
    activ = alpha_hat!= 0
    
    weighting = (1 / freq_alphahat)^(1/2)
    weighting[weighting==Inf] = 0
    weighting[weighting > 5] = 5
    unique = as.numeric((alpha_hat[activ] * weighting[activ]) %*% geno[activ,])
    new_ebv = (unique - mean(unique))/sd(unique)
    
    population = insert.bve(population, bves = cbind(names(ebv), new_ebv))
    population = breeding.diploid(population, selection.size = sel_size, breeding.size = pop_size,
                                  selection.criteria = "bve",
                                  share.genotyped = 1,
                                  display.progress = display.progress)
    
  } else if(scenario == 189){
    
    activ = alpha_hat!= 0
    
    weighting = (1 / freq_alphahat)^(1/3)
    weighting[weighting==Inf] = 0
    
    unique = as.numeric((alpha_hat[activ] * weighting[activ]) %*% geno[activ,])
    new_ebv = (unique - mean(unique))/sd(unique)
    
    population = insert.bve(population, bves = cbind(names(ebv), new_ebv))
    population = breeding.diploid(population, selection.size = sel_size, breeding.size = pop_size,
                                  selection.criteria = "bve",
                                  share.genotyped = 1,
                                  display.progress = display.progress)
    
  } else if(scenario == 190){
    
    activ = alpha_hat!= 0
    
    weighting = (1 / freq_alphahat)^(1/4)
    weighting[weighting==Inf] = 0
    
    unique = as.numeric((alpha_hat[activ] * weighting[activ]) %*% geno[activ,])
    new_ebv = (unique - mean(unique))/sd(unique)
    
    population = insert.bve(population, bves = cbind(names(ebv), new_ebv))
    population = breeding.diploid(population, selection.size = sel_size, breeding.size = pop_size,
                                  selection.criteria = "bve",
                                  share.genotyped = 1,
                                  display.progress = display.progress)
    
  } else if(scenario == 191){
    
    activ = alpha_hat!= 0
    
    weighting = 0.8 + 0.2 * (1 / freq_alphahat)^(1/2)
    weighting[weighting==Inf] = 0
    
    unique = as.numeric((alpha_hat[activ] * weighting[activ]) %*% geno[activ,])
    new_ebv = (unique - mean(unique))/sd(unique)
    
    population = insert.bve(population, bves = cbind(names(ebv), new_ebv))
    population = breeding.diploid(population, selection.size = sel_size, breeding.size = pop_size,
                                  selection.criteria = "bve",
                                  share.genotyped = 1,
                                  display.progress = display.progress)
    
  } else if(scenario == 192){
    
    activ = alpha_hat!= 0
    
    weighting = 0.6 + 0.4 * (1 / freq_alphahat)^(1/2)
    weighting[weighting==Inf] = 0
    
    unique = as.numeric((alpha_hat[activ] * weighting[activ]) %*% geno[activ,])
    new_ebv = (unique - mean(unique))/sd(unique)
    
    population = insert.bve(population, bves = cbind(names(ebv), new_ebv))
    population = breeding.diploid(population, selection.size = sel_size, breeding.size = pop_size,
                                  selection.criteria = "bve",
                                  share.genotyped = 1,
                                  display.progress = display.progress)
    
  } else if(scenario == 193){
    
    activ = alpha_hat!= 0
    
    weighting = (pi/2 - asin(sqrt(freq_alphahat)))/(sqrt(freq_alphahat * (1-freq_alphahat)))
    weighting[weighting==Inf] = 0
    
    unique = as.numeric((alpha_hat[activ] * weighting[activ]) %*% geno[activ,])
    new_ebv = (unique - mean(unique))/sd(unique)
    
    population = insert.bve(population, bves = cbind(names(ebv), new_ebv))
    population = breeding.diploid(population, selection.size = sel_size, breeding.size = pop_size,
                                  selection.criteria = "bve",
                                  share.genotyped = 1,
                                  display.progress = display.progress)
    
  } else if( scenario == 231){
    

    work_dir = paste0("mixblup_", scenario, "_", seed)
    if(index == 1){
      dir.create(work_dir)
      system(paste0("cp /home/WUR/pook001/SysDir.inp ", work_dir, "/SysDir.inp"))
      setwd(work_dir)
      
      system(paste0("cp /home/WUR/pook001/msv msv"))
      system("chmod +x msv")
    }
    
    get.plink(population, gen =bve.gen,
              path = "paternal_plink", fam.id = TRUE, type = 1)
    
    get.plink(population, gen =bve.gen,
              path = "maternal_plink", fam.id = TRUE, type = 2)
    
    write.table(file = "EBV.txt", cbind(get.id(population, gen = bve.gen), ebv), row.names = FALSE, col.names = FALSE, quote = FALSE)
    write.table(file = "sol.dat", cbind(1,1,1,1:length(alpha_hat), alpha_hat), row.names = FALSE, col.names = FALSE, quote = FALSE)
    write.table(file = "all_freq", cbind(1:length(alpha_hat), 0), row.names = FALSE, col.names = FALSE, quote = FALSE)
    
    system("./msv --matgeno maternal_plink.bed --patgeno paternal_plink.bed --freq all_freq --ebv EBV.txt --sol sol.dat --snpeff 1 --nrgen 2 --selprop 0.1 --bp2M 1e-08")
    
    a = read.table("EBV.txt")
    b = read.table("genetic_MSVs_1gen.txt")
    c = read.table("Index_5_6.txt") #Piter Index 5/6
    d = read.table("ExpBVSelGrOff_per_individual.txt") # Tobias Index
    
    new_ebv = d[,2]
    
    insert.bve = cbind(names(ebv), new_ebv)
    population = insert.bve(population, bves = insert.bve )
    
    acc2 = rbind(acc2, t(analyze.bv(population, gen = bve.gen)[[1]]))
    
    pop1 = breeding.diploid(population, bve = TRUE, bve.gen = bve.gen)
    
    acc1 = rbind(acc1, t(analyze.bv(pop1, gen = bve.gen)[[1]]))
    
    print(colMeans(acc1))
    print(colMeans(acc2))
    
    # Selection based on genomic breeding value
    population = breeding.diploid(population, selection.size = sel_size, breeding.size = pop_size,
                                  selection.criteria = "bve",
                                  share.genotyped = 1,
                                  display.progress = display.progress)
    
    
    
  } else if( scenario == 233){
    
    
    work_dir = paste0("mixblup_", scenario, "_", seed)
    if(index == 1){
      dir.create(work_dir)
      system(paste0("cp /home/WUR/pook001/SysDir.inp ", work_dir, "/SysDir.inp"))
      setwd(work_dir)
      
      system(paste0("cp /home/WUR/pook001/msv msv"))
      system("chmod +x msv")
    }
    
    get.plink(population, gen =bve.gen,
              path = "paternal_plink", fam.id = TRUE, type = 1)
    
    get.plink(population, gen =bve.gen,
              path = "maternal_plink", fam.id = TRUE, type = 2)
    
    write.table(file = "EBV.txt", cbind(get.id(population, gen = bve.gen), ebv), row.names = FALSE, col.names = FALSE, quote = FALSE)
    write.table(file = "sol.dat", cbind(1,1,1,1:length(alpha_hat), alpha_hat), row.names = FALSE, col.names = FALSE, quote = FALSE)
    write.table(file = "all_freq", cbind(1:length(alpha_hat), 0), row.names = FALSE, col.names = FALSE, quote = FALSE)
    
    system("./msv --matgeno maternal_plink.bed --patgeno paternal_plink.bed --freq all_freq --ebv EBV.txt --sol sol.dat --snpeff 1 --nrgen 2 --selprop 0.1 --bp2M 1e-08")
    
    a = read.table("EBV.txt")
    b = read.table("genetic_MSVs_1gen.txt")
    c = read.table("Index_5_6.txt") #Piter Index 5/6
    d = read.table("ExpBVSelGrOff_per_individual.txt") # Tobias Index
    
    new_ebv = c[,2]
    
    insert.bve = cbind(names(ebv), new_ebv)
    population = insert.bve(population, bves = insert.bve )
    
    acc2 = rbind(acc2, t(analyze.bv(population, gen = bve.gen)[[1]]))
    
    pop1 = breeding.diploid(population, bve = TRUE, bve.gen = bve.gen)
    
    acc1 = rbind(acc1, t(analyze.bv(pop1, gen = bve.gen)[[1]]))
    
    print(colMeans(acc1))
    print(colMeans(acc2))
    
    # Selection based on genomic breeding value
    population = breeding.diploid(population, selection.size = sel_size, breeding.size = pop_size,
                                  selection.criteria = "bve",
                                  share.genotyped = 1,
                                  display.progress = display.progress)
    
    
    
  } else if( scenario == 234){
    
    
    work_dir = paste0("mixblup_", scenario, "_", seed)
    if(index == 1){
      dir.create(work_dir)
      system(paste0("cp /home/WUR/pook001/SysDir.inp ", work_dir, "/SysDir.inp"))
      setwd(work_dir)
      
      system(paste0("cp /home/WUR/pook001/msv msv"))
      system("chmod +x msv")
    }
    
    get.plink(population, gen =bve.gen,
              path = "paternal_plink", fam.id = TRUE, type = 1)
    
    get.plink(population, gen =bve.gen,
              path = "maternal_plink", fam.id = TRUE, type = 2)
    
    write.table(file = "EBV.txt", cbind(get.id(population, gen = bve.gen), ebv), row.names = FALSE, col.names = FALSE, quote = FALSE)
    write.table(file = "sol.dat", cbind(1,1,1,1:length(alpha_hat), alpha_hat), row.names = FALSE, col.names = FALSE, quote = FALSE)
    write.table(file = "all_freq", cbind(1:length(alpha_hat), 0), row.names = FALSE, col.names = FALSE, quote = FALSE)
    
    system("./msv --matgeno maternal_plink.bed --patgeno paternal_plink.bed --freq all_freq --ebv EBV.txt --sol sol.dat --snpeff 1 --nrgen 2 --selprop 0.1 --bp2M 1e-08")
    
    a = read.table("EBV.txt")
    b = read.table("genetic_MSVs_1gen.txt")
    c = read.table("Index_5_6.txt") #Piter Index 5/6
    d = read.table("ExpBVSelGrOff_per_individual.txt") # Tobias Index
    
    new_ebv = c[,3]
    
    insert.bve = cbind(names(ebv), new_ebv)
    population = insert.bve(population, bves = insert.bve )
    
    acc2 = rbind(acc2, t(analyze.bv(population, gen = bve.gen)[[1]]))
    
    pop1 = breeding.diploid(population, bve = TRUE, bve.gen = bve.gen)
    
    acc1 = rbind(acc1, t(analyze.bv(pop1, gen = bve.gen)[[1]]))
    
    print(colMeans(acc1))
    print(colMeans(acc2))
    
    # Selection based on genomic breeding value
    population = breeding.diploid(population, selection.size = sel_size, breeding.size = pop_size,
                                  selection.criteria = "bve",
                                  share.genotyped = 1,
                                  display.progress = display.progress)
    
    
    
  } else if( scenario == 237){
    
    targetdegree = 5
    temp_geno <- geno
    pi1 <- rowMeans(temp_geno)/2
    temp_kin <- crossprod(temp_geno- 2*pi1) / sum(pi1*(1-pi1)) / 2
    
    selection_index <- as.numeric(ebv)
    
    diag_kin <- diag(temp_kin)
    kin_mat <- temp_kin - min(temp_kin)
    kin_mat <- kin_mat/max(as.vector(kin_mat))*2
    kin_mat_tri <- kin_mat[upper.tri(kin_mat)]
    kin_mat <- t(kin_mat)
    kin_mat[upper.tri(kin_mat)] <- kin_mat_tri
    
    dir.create(paste0("alphamaterun/unique_", targetdegree, "_", scenario,"_", seed))
    
    write.table(kin_mat, sprintf("./alphamaterun/unique_%i_%i_%i/Nrm.txt", targetdegree,scenario,seed), sep="\t", row.names=TRUE, col.names=FALSE)
    
    criterion <- data.frame(Genotype = names(ebv),
                            Value = as.vector(ebv))
    write.table(criterion, sprintf("./alphamaterun/unique_%i_%i_%i/Criterion.txt", targetdegree,scenario,seed), sep="\t", row.names=FALSE, quote=FALSE, col.names=FALSE)
    
    gender_file = cbind(names(ebv), c(rep(1, pop_size[1]), rep(2, pop_size[2])))
    write.table(gender_file, sprintf("./alphamaterun/unique_%i_%i_%i/Gender.txt", targetdegree,scenario,seed), sep="\t", row.names=FALSE, col.names=FALSE, quote = FALSE)
    
    spec_text <- sprintf("Seed , %i\nNrmMatrixFile , Nrm.txt\nGenderFile , Gender.txt\nSelCriterionFile , Criterion.txt\nNumberOfMatings , %i\nNumberOfMaleParents , %i\nNumberOfFemaleParents , %i\nTargetDegree , %i\nEqualizeMaleContributions , Yes\nEqualizeFemaleContributions , Yes\nPreselectMales , Yes\nPreselectMalePercentage , 100\nPreselectFemales , Yes\nPreselectFemalePercentage , 100\nEvolAlgNumberOfIterations , 1000\nEvolAlgNumberOfIterationsPrint , 5\nEvolAlgNumberOfIterationsStop , 50\nStop",
                         seed, sum(pop_size), sel_size[1], sel_size[2],targetdegree)
    
    write(x = spec_text, file=sprintf("./alphamaterun/unique_%i_%i_%i/AlphaMateSpec.txt", targetdegree, scenario,seed))
    
    sim_path = getwd()
    setwd(paste0("alphamaterun/unique_", targetdegree, "_", scenario,"_", seed))
    
    
    
    suppressWarnings(file.remove("MatingPlanModeOptTarget1.txt"))
    
    if(seed<3){
      system(sprintf(paste0( sim_path, "/AlphaMate/binaries/AlphaMate_Unix")))
      
    } else{
      system(sprintf(paste0( sim_path, "/AlphaMate_0.2.0/bin/AlphaMate_Linux_0.2.02")))
      
    }
    
    setwd(sim_path)
    
    matings <- read.table(sprintf("./alphamaterun/unique_%i_%i_%i/MatingPlanModeOptTarget1.txt", targetdegree, scenario, seed), sep="", header=TRUE)
    
    males = match(as.character(matings$Parent1), names(ebv)[1:pop_size[1]])
    females = match(as.character(matings$Parent2), names(ebv)[-(1:pop_size[1])])
    
    sex = rep(0, sum(pop_size))
    sex[sample(sum(pop_size), pop_size[2])] = 1
    fixed_breeding = cbind(get.ngen(population), 1, males, get.ngen(population),2,females, sex)
    # Selection based on genomic breeding value
    population = breeding.diploid(population,
                                  fixed.breeding = fixed_breeding,
                                  share.genotyped = 1,
                                  display.progress = display.progress)
    
    
  }else if( scenario == 238){
    
    targetdegree = 15
    temp_geno <- geno
    pi1 <- rowMeans(temp_geno)/2
    temp_kin <- crossprod(temp_geno- 2*pi1) / sum(pi1*(1-pi1)) / 2
    
    selection_index <- as.numeric(ebv)
    
    diag_kin <- diag(temp_kin)
    kin_mat <- temp_kin - min(temp_kin)
    kin_mat <- kin_mat/max(as.vector(kin_mat))*2
    kin_mat_tri <- kin_mat[upper.tri(kin_mat)]
    kin_mat <- t(kin_mat)
    kin_mat[upper.tri(kin_mat)] <- kin_mat_tri
    
    dir.create(paste0("alphamaterun/unique_", targetdegree, "_", scenario,"_", seed))
    
    write.table(kin_mat, sprintf("./alphamaterun/unique_%i_%i_%i/Nrm.txt", targetdegree,scenario,seed), sep="\t", row.names=TRUE, col.names=FALSE)
    
    criterion <- data.frame(Genotype = names(ebv),
                            Value = as.vector(ebv))
    write.table(criterion, sprintf("./alphamaterun/unique_%i_%i_%i/Criterion.txt", targetdegree,scenario,seed), sep="\t", row.names=FALSE, quote=FALSE, col.names=FALSE)
    
    gender_file = cbind(names(ebv), c(rep(1, pop_size[1]), rep(2, pop_size[2])))
    write.table(gender_file, sprintf("./alphamaterun/unique_%i_%i_%i/Gender.txt", targetdegree,scenario,seed), sep="\t", row.names=FALSE, col.names=FALSE, quote = FALSE)
    
    spec_text <- sprintf("Seed , %i\nNrmMatrixFile , Nrm.txt\nGenderFile , Gender.txt\nSelCriterionFile , Criterion.txt\nNumberOfMatings , %i\nNumberOfMaleParents , %i\nNumberOfFemaleParents , %i\nTargetDegree , %i\nEqualizeMaleContributions , Yes\nEqualizeFemaleContributions , Yes\nPreselectMales , Yes\nPreselectMalePercentage , 50\nPreselectFemales , Yes\nPreselectFemalePercentage , 50\nEvolAlgNumberOfIterations , 1000\nEvolAlgNumberOfIterationsPrint , 5\nEvolAlgNumberOfIterationsStop , 50\nStop",
                         seed, sum(pop_size), sel_size[1], sel_size[2],targetdegree)
    
    write(x = spec_text, file=sprintf("./alphamaterun/unique_%i_%i_%i/AlphaMateSpec.txt", targetdegree, scenario,seed))
    
    sim_path = getwd()
    setwd(paste0("alphamaterun/unique_", targetdegree, "_", scenario,"_", seed))
    
    
    
    suppressWarnings(file.remove("MatingPlanModeOptTarget1.txt"))
    
    if(seed<3){
      system(sprintf(paste0( sim_path, "/AlphaMate/binaries/AlphaMate_Unix")))
      
    } else{
      system(sprintf(paste0( sim_path, "/AlphaMate_0.2.0/bin/AlphaMate_Linux_0.2.02")))
      
    }
    
    setwd(sim_path)
    
    matings <- read.table(sprintf("./alphamaterun/unique_%i_%i_%i/MatingPlanModeOptTarget1.txt", targetdegree, scenario, seed), sep="", header=TRUE)
    
    males = match(as.character(matings$Parent1), names(ebv)[1:pop_size[1]])
    females = match(as.character(matings$Parent2), names(ebv)[-(1:pop_size[1])])
    
    sex = rep(0, sum(pop_size))
    sex[sample(sum(pop_size), pop_size[2])] = 1
    fixed_breeding = cbind(get.ngen(population), 1, males, get.ngen(population),2,females, sex)
    # Selection based on genomic breeding value
    population = breeding.diploid(population,
                                  fixed.breeding = fixed_breeding,
                                  share.genotyped = 1,
                                  display.progress = display.progress)
    
    
  }else if( scenario == 239){
    targetdegree = 30
    temp_geno <- geno
    pi1 <- rowMeans(temp_geno)/2
    temp_kin <- crossprod(temp_geno- 2*pi1) / sum(pi1*(1-pi1)) / 2
    
    selection_index <- as.numeric(ebv)
    
    diag_kin <- diag(temp_kin)
    kin_mat <- temp_kin - min(temp_kin)
    kin_mat <- kin_mat/max(as.vector(kin_mat))*2
    kin_mat_tri <- kin_mat[upper.tri(kin_mat)]
    kin_mat <- t(kin_mat)
    kin_mat[upper.tri(kin_mat)] <- kin_mat_tri
    
    dir.create(paste0("alphamaterun/unique_", targetdegree, "_", scenario,"_", seed))
    
    write.table(kin_mat, sprintf("./alphamaterun/unique_%i_%i_%i/Nrm.txt", targetdegree,scenario,seed), sep="\t", row.names=TRUE, col.names=FALSE)
    
    criterion <- data.frame(Genotype = names(ebv),
                            Value = as.vector(ebv))
    write.table(criterion, sprintf("./alphamaterun/unique_%i_%i_%i/Criterion.txt", targetdegree,scenario,seed), sep="\t", row.names=FALSE, quote=FALSE, col.names=FALSE)
    
    gender_file = cbind(names(ebv), c(rep(1, pop_size[1]), rep(2, pop_size[2])))
    write.table(gender_file, sprintf("./alphamaterun/unique_%i_%i_%i/Gender.txt", targetdegree,scenario,seed), sep="\t", row.names=FALSE, col.names=FALSE, quote = FALSE)
    
    spec_text <- sprintf("Seed , %i\nNrmMatrixFile , Nrm.txt\nGenderFile , Gender.txt\nSelCriterionFile , Criterion.txt\nNumberOfMatings , %i\nNumberOfMaleParents , %i\nNumberOfFemaleParents , %i\nTargetDegree , %i\nEqualizeMaleContributions , Yes\nEqualizeFemaleContributions , Yes\nPreselectMales , Yes\nPreselectMalePercentage , 33\nPreselectFemales , Yes\nPreselectFemalePercentage , 33\nEvolAlgNumberOfIterations , 1000\nEvolAlgNumberOfIterationsPrint , 5\nEvolAlgNumberOfIterationsStop , 50\nStop",
                         seed, sum(pop_size), sel_size[1], sel_size[2],targetdegree)
    
    write(x = spec_text, file=sprintf("./alphamaterun/unique_%i_%i_%i/AlphaMateSpec.txt", targetdegree, scenario,seed))
    
    sim_path = getwd()
    setwd(paste0("alphamaterun/unique_", targetdegree, "_", scenario,"_", seed))
    
    
    
    suppressWarnings(file.remove("MatingPlanModeOptTarget1.txt"))
    
    if(seed<3){
      system(sprintf(paste0( sim_path, "/AlphaMate/binaries/AlphaMate_Unix")))
      
    } else{
      system(sprintf(paste0( sim_path, "/AlphaMate_0.2.0/bin/AlphaMate_Linux_0.2.02")))
      
    }
    
    setwd(sim_path)
    
    matings <- read.table(sprintf("./alphamaterun/unique_%i_%i_%i/MatingPlanModeOptTarget1.txt", targetdegree, scenario, seed), sep="", header=TRUE)
    
    males = match(as.character(matings$Parent1), names(ebv)[1:pop_size[1]])
    females = match(as.character(matings$Parent2), names(ebv)[-(1:pop_size[1])])
    
    sex = rep(0, sum(pop_size))
    sex[sample(sum(pop_size), pop_size[2])] = 1
    fixed_breeding = cbind(get.ngen(population), 1, males, get.ngen(population),2,females, sex)
    # Selection based on genomic breeding value
    population = breeding.diploid(population,
                                  fixed.breeding = fixed_breeding,
                                  share.genotyped = 1,
                                  display.progress = display.progress)
    
    
  }else if( scenario == 240){
    
    targetdegree = 45
    temp_geno <- geno
    pi1 <- rowMeans(temp_geno)/2
    temp_kin <- crossprod(temp_geno- 2*pi1) / sum(pi1*(1-pi1)) / 2
    
    selection_index <- as.numeric(ebv)
    
    diag_kin <- diag(temp_kin)
    kin_mat <- temp_kin - min(temp_kin)
    kin_mat <- kin_mat/max(as.vector(kin_mat))*2
    kin_mat_tri <- kin_mat[upper.tri(kin_mat)]
    kin_mat <- t(kin_mat)
    kin_mat[upper.tri(kin_mat)] <- kin_mat_tri
    
    dir.create(paste0("alphamaterun/unique_", targetdegree, "_", scenario,"_", seed))
    
    write.table(kin_mat, sprintf("./alphamaterun/unique_%i_%i_%i/Nrm.txt", targetdegree,scenario,seed), sep="\t", row.names=TRUE, col.names=FALSE)
    
    criterion <- data.frame(Genotype = names(ebv),
                            Value = as.vector(ebv))
    write.table(criterion, sprintf("./alphamaterun/unique_%i_%i_%i/Criterion.txt", targetdegree,scenario,seed), sep="\t", row.names=FALSE, quote=FALSE, col.names=FALSE)
    
    gender_file = cbind(names(ebv), c(rep(1, pop_size[1]), rep(2, pop_size[2])))
    write.table(gender_file, sprintf("./alphamaterun/unique_%i_%i_%i/Gender.txt", targetdegree,scenario,seed), sep="\t", row.names=FALSE, col.names=FALSE, quote = FALSE)
    
    spec_text <- sprintf("Seed , %i\nNrmMatrixFile , Nrm.txt\nGenderFile , Gender.txt\nSelCriterionFile , Criterion.txt\nNumberOfMatings , %i\nNumberOfMaleParents , %i\nNumberOfFemaleParents , %i\nTargetDegree , %i\nEqualizeMaleContributions , Yes\nEqualizeFemaleContributions , Yes\nPreselectMales , Yes\nPreselectMalePercentage , 20\nPreselectFemales , Yes\nPreselectFemalePercentage , 20\nEvolAlgNumberOfIterations , 1000\nEvolAlgNumberOfIterationsPrint , 5\nEvolAlgNumberOfIterationsStop , 50\nStop",
                         seed, sum(pop_size), sel_size[1], sel_size[2],targetdegree)
    
    write(x = spec_text, file=sprintf("./alphamaterun/unique_%i_%i_%i/AlphaMateSpec.txt", targetdegree, scenario,seed))
    
    sim_path = getwd()
    setwd(paste0("alphamaterun/unique_", targetdegree, "_", scenario,"_", seed))
    
    
    
    suppressWarnings(file.remove("MatingPlanModeOptTarget1.txt"))
    
    if(seed<3){
      system(sprintf(paste0( sim_path, "/AlphaMate/binaries/AlphaMate_Unix")))
      
    } else{
      system(sprintf(paste0( sim_path, "/AlphaMate_0.2.0/bin/AlphaMate_Linux_0.2.02")))
      
    }
    
    setwd(sim_path)
    
    matings <- read.table(sprintf("./alphamaterun/unique_%i_%i_%i/MatingPlanModeOptTarget1.txt", targetdegree, scenario, seed), sep="", header=TRUE)
    
    males = match(as.character(matings$Parent1), names(ebv)[1:pop_size[1]])
    females = match(as.character(matings$Parent2), names(ebv)[-(1:pop_size[1])])
    
    sex = rep(0, sum(pop_size))
    sex[sample(sum(pop_size), pop_size[2])] = 1
    fixed_breeding = cbind(get.ngen(population), 1, males, get.ngen(population),2,females, sex)
    # Selection based on genomic breeding value
    population = breeding.diploid(population,
                                  fixed.breeding = fixed_breeding,
                                  share.genotyped = 1,
                                  display.progress = display.progress)
    
    
  } else if( scenario == 247){
    
    # Selection based on genomic breeding value
    population = breeding.diploid(population, selection.size = c(50,125), breeding.size = pop_size,
                                  selection.criteria = "bve",
                                  share.genotyped = 1,
                                  display.progress = display.progress)
    
  } else if( scenario == 248){
    
    # Selection based on genomic breeding value
    population = breeding.diploid(population, selection.size = c(45,112), breeding.size = pop_size,
                                  selection.criteria = "bve",
                                  share.genotyped = 1,
                                  display.progress = display.progress)
    
  } else if( scenario == 249){
    
    # Selection based on genomic breeding value
    population = breeding.diploid(population, selection.size = c(42,105), breeding.size = pop_size,
                                  selection.criteria = "bve",
                                  share.genotyped = 1,
                                  display.progress = display.progress)
    
  } else if( scenario == 250){
    
    # Selection based on genomic breeding value
    population = breeding.diploid(population, selection.size = c(38,95), breeding.size = pop_size,
                                  selection.criteria = "bve",
                                  share.genotyped = 1,
                                  display.progress = display.progress)
    
  } else if( scenario == 251){
    
    # Selection based on genomic breeding value
    population = breeding.diploid(population, selection.size = c(36,90), breeding.size = pop_size,
                                  selection.criteria = "bve",
                                  share.genotyped = 1,
                                  display.progress = display.progress)
    
  } else if( scenario == 252){
    
    # Selection based on genomic breeding value
    population = breeding.diploid(population, selection.size = c(34,85), breeding.size = pop_size,
                                  selection.criteria = "bve",
                                  share.genotyped = 1,
                                  display.progress = display.progress)
    
  } else if( scenario == 253){
    
    # Selection based on genomic breeding value
    population = breeding.diploid(population, selection.size = c(32,80), breeding.size = pop_size,
                                  selection.criteria = "bve",
                                  share.genotyped = 1,
                                  display.progress = display.progress)
    
  } else if( scenario == 254){
    
    # Selection based on genomic breeding value
    population = breeding.diploid(population, selection.size = c(30,75), breeding.size = pop_size,
                                  selection.criteria = "bve",
                                  share.genotyped = 1,
                                  display.progress = display.progress)
    
  } else if( scenario == 255){
    
    # Selection based on genomic breeding value
    population = breeding.diploid(population, selection.size = c(28,70), breeding.size = pop_size,
                                  selection.criteria = "bve",
                                  share.genotyped = 1,
                                  display.progress = display.progress)
    
  } else if(scenario == 303){
    
    weight_rare1 = 1.0
    weight_rare2 = 0.0
    avoid_mating_inb = 0.5
    avoid_mating_kin = 0.95
    weight_kinship1 = 0.12
    weight_kinship2 = 0.02
    ogc_scaler = 1.5
    ogc = TRUE
    
    sel_size = round(c(40, 100) * 0.32)
    
    activ = alpha_hat!= 0
    
    weighting = (1 / freq_alphahat)^(1/3)
    weighting[weighting==Inf] = 0
    weighting[weighting>10] = 10
    
    bve_rare =  as.numeric((alpha_hat[activ] * weighting[activ]) %*% geno[activ,])
    bve_rare1 = (bve_rare-mean(bve_rare)) /sd(bve_rare)
    
    weighting = (1 / freq_alphahat)^(1/2)
    weighting[weighting==Inf] = 0
    weighting[weighting>10] = 10
    
    bve_rare =  as.numeric((alpha_hat[activ] * weighting[activ]) %*% geno[activ,])
    bve_rare2 = (bve_rare-mean(bve_rare)) /sd(bve_rare)
    
    cand = breeding.diploid(population, selection.size = sel_size, breeding.size = pop_size,
                            selection.criteria = "bve", export.selected = TRUE,
                            verbose = FALSE)
    
    A = kinship.exp(population, gen = get.ngen(population))
    diag(A) = (diag(A)-0.5)*2
    
    rel_top = colMeans(A[cand[[1]][,3],]) + colMeans(A[cand[[2]][,3] + pop_size[1],])
    unique = -rel_top #
    bve_kinship1 = (unique - mean(unique))/sd(unique)
    
    rel_top = colMeans(A)
    unique = -rel_top #
    bve_kinship2 = (unique - mean(unique))/sd(unique)
    
    bve_plain = (ebv - mean(ebv))/sd(ebv)
    
    
    new_ebv = weight_rare1 * bve_rare1 + weight_rare2 * bve_rare2 + weight_kinship1 * bve_kinship1 + weight_kinship2 * bve_kinship2  + (1-weight_rare1 - weight_rare2 - weight_kinship1 - weight_kinship2) * bve_plain
    
    to_insert = cbind(names(ebv), new_ebv)
    
    population = insert.bve(population, bves = to_insert)
    
    sel_size_tmp = sel_size
    sel_size_tmp[1] = pop_size[1]
    
    population = breeding.diploid(population, selection.size = sel_size_tmp, breeding.size = pop_size,
                                  selection.criteria = "bve",
                                  share.genotyped = 1,
                                  avoid.mating.inb.quantile = avoid_mating_inb,
                                  avoid.mating.kinship.quantile = avoid_mating_kin,
                                  avoid.mating.kinship.gen = get.ngen(population),
                                  display.progress = display.progress,
                                  ogc = ogc,
                                  ogc.target = "max.BV",
                                  ogc.uniform = "female",
                                  ogc.ub.sKin.increase = 0.005 * ogc_scaler,
                                  verbose = TRUE)
  } else if(scenario == 304){
    
    weight_rare1 = 0.4030
    weight_rare2 = 0.0020
    avoid_mating_inb = 0.900
    avoid_mating_kin = 1.000
    weight_kinship1 = 0.2030
    weight_kinship2 = 0.0290
    ogc_scaler = 1.000
    ogc = TRUE
    
    sel_size = round(c(40, 100) * 0.83)
    
    activ = alpha_hat!= 0
    
    weighting = (1 / freq_alphahat)^(1/3)
    weighting[weighting==Inf] = 0
    weighting[weighting>10] = 10
    
    bve_rare =  as.numeric((alpha_hat[activ] * weighting[activ]) %*% geno[activ,])
    bve_rare1 = (bve_rare-mean(bve_rare)) /sd(bve_rare)
    
    weighting = (1 / freq_alphahat)^(1/2)
    weighting[weighting==Inf] = 0
    weighting[weighting>10] = 10
    
    bve_rare =  as.numeric((alpha_hat[activ] * weighting[activ]) %*% geno[activ,])
    bve_rare2 = (bve_rare-mean(bve_rare)) /sd(bve_rare)
    
    cand = breeding.diploid(population, selection.size = sel_size, breeding.size = pop_size,
                            selection.criteria = "bve", export.selected = TRUE,
                            verbose = FALSE)
    
    A = kinship.exp(population, gen = get.ngen(population))
    diag(A) = (diag(A)-0.5)*2
    
    rel_top = colMeans(A[cand[[1]][,3],]) + colMeans(A[cand[[2]][,3] + pop_size[1],])
    unique = -rel_top #
    bve_kinship1 = (unique - mean(unique))/sd(unique)
    
    rel_top = colMeans(A)
    unique = -rel_top #
    bve_kinship2 = (unique - mean(unique))/sd(unique)
    
    bve_plain = (ebv - mean(ebv))/sd(ebv)
    
    
    new_ebv = weight_rare1 * bve_rare1 + weight_rare2 * bve_rare2 + weight_kinship1 * bve_kinship1 + weight_kinship2 * bve_kinship2  + (1-weight_rare1 - weight_rare2 - weight_kinship1 - weight_kinship2) * bve_plain
    
    to_insert = cbind(names(ebv), new_ebv)
    
    population = insert.bve(population, bves = to_insert)
    
    sel_size_tmp = sel_size
    sel_size_tmp[1] = pop_size[1]
    
    population = breeding.diploid(population, selection.size = sel_size_tmp, breeding.size = pop_size,
                                  selection.criteria = "bve",
                                  share.genotyped = 1,
                                  avoid.mating.inb.quantile = avoid_mating_inb,
                                  avoid.mating.kinship.quantile = avoid_mating_kin,
                                  avoid.mating.kinship.gen = get.ngen(population),
                                  display.progress = display.progress,
                                  ogc = ogc,
                                  ogc.target = "max.BV",
                                  ogc.uniform = "female",
                                  ogc.ub.sKin.increase = 0.005 * ogc_scaler,
                                  verbose = TRUE)
  } else if(scenario == 320){
    
    activ = alpha_real!= 0
    
    weighting = (1 / freq_alpha)^(1/2)
    weighting[weighting==Inf] = 0
    
    unique = as.numeric((alpha_real[activ] * weighting[activ]) %*% geno[activ,])
    new_ebv = (unique - mean(unique))/sd(unique)
    
    population = insert.bve(population, bves = cbind(names(ebv), new_ebv))
    population = breeding.diploid(population, selection.size = sel_size, breeding.size = pop_size,
                                  selection.criteria = "bve",
                                  share.genotyped = 1,
                                  display.progress = display.progress)
    
  } else if(scenario == 321){
    
    activ = alpha_real!= 0
    
    weighting = (1 / freq_alpha)^(1/2)
    weighting[weighting==Inf] = 0
    weighting[weighting > 5] = 5
    unique = as.numeric((alpha_real[activ] * weighting[activ]) %*% geno[activ,])
    new_ebv = (unique - mean(unique))/sd(unique)
    
    population = insert.bve(population, bves = cbind(names(ebv), new_ebv))
    population = breeding.diploid(population, selection.size = sel_size, breeding.size = pop_size,
                                  selection.criteria = "bve",
                                  share.genotyped = 1,
                                  display.progress = display.progress)
    
  } else if(scenario == 322){
    
    activ = alpha_real!= 0
    
    weighting = (1 / freq_alpha)^(1/3)
    weighting[weighting==Inf] = 0
    
    unique = as.numeric((alpha_real[activ] * weighting[activ]) %*% geno[activ,])
    new_ebv = (unique - mean(unique))/sd(unique)
    
    population = insert.bve(population, bves = cbind(names(ebv), new_ebv))
    population = breeding.diploid(population, selection.size = sel_size, breeding.size = pop_size,
                                  selection.criteria = "bve",
                                  share.genotyped = 1,
                                  display.progress = display.progress)
    
  } else if(scenario == 323){
    
    activ = alpha_real!= 0
    
    weighting = (1 / freq_alpha)^(1/4)
    weighting[weighting==Inf] = 0
    
    unique = as.numeric((alpha_real[activ] * weighting[activ]) %*% geno[activ,])
    new_ebv = (unique - mean(unique))/sd(unique)
    
    population = insert.bve(population, bves = cbind(names(ebv), new_ebv))
    population = breeding.diploid(population, selection.size = sel_size, breeding.size = pop_size,
                                  selection.criteria = "bve",
                                  share.genotyped = 1,
                                  display.progress = display.progress)
    
  } else if(scenario == 324){
    
    activ = alpha_real!= 0
    
    weighting = 0.8 + 0.2 * (1 / freq_alpha)^(1/2)
    weighting[weighting==Inf] = 0
    
    unique = as.numeric((alpha_real[activ] * weighting[activ]) %*% geno[activ,])
    new_ebv = (unique - mean(unique))/sd(unique)
    
    population = insert.bve(population, bves = cbind(names(ebv), new_ebv))
    population = breeding.diploid(population, selection.size = sel_size, breeding.size = pop_size,
                                  selection.criteria = "bve",
                                  share.genotyped = 1,
                                  display.progress = display.progress)
    
  } else if(scenario == 325){
    
    activ = alpha_real!= 0
    
    weighting = 0.6 + 0.4 * (1 / freq_alpha)^(1/2)
    weighting[weighting==Inf] = 0
    
    unique = as.numeric((alpha_real[activ] * weighting[activ]) %*% geno[activ,])
    new_ebv = (unique - mean(unique))/sd(unique)
    
    population = insert.bve(population, bves = cbind(names(ebv), new_ebv))
    
    population = breeding.diploid(population, selection.size = sel_size, breeding.size = pop_size,
                                  selection.criteria = "bve",
                                  share.genotyped = 1,
                                  display.progress = display.progress)
    
  } else if(scenario == 326){
    
    activ = alpha_real!= 0
    
    weighting = (pi/2 - asin(sqrt(freq_alpha)))/(sqrt(freq_alpha * (1-freq_alpha)))
    weighting[weighting==Inf] = 0
    
    unique = as.numeric((alpha_real[activ] * weighting[activ]) %*% geno[activ,])
    new_ebv = (unique - mean(unique))/sd(unique)
    
    population = insert.bve(population, bves = cbind(names(ebv), new_ebv))
    population = breeding.diploid(population, selection.size = sel_size, breeding.size = pop_size,
                                  selection.criteria = "bve",
                                  share.genotyped = 1,
                                  display.progress = display.progress)
    
  } else if( scenario == 335){
    
    
    work_dir = paste0("mixblup_", scenario, "_", seed)
    if(index == 1){
      dir.create(work_dir)
      system(paste0("cp /home/WUR/pook001/SysDir.inp ", work_dir, "/SysDir.inp"))
      setwd(work_dir)
      
      system(paste0("cp /home/WUR/pook001/msv msv"))
      system("chmod +x msv")
    }
    
    get.plink(population, gen =bve.gen1,
              path = "paternal_plink", fam.id = TRUE, type = 1)
    
    get.plink(population, gen =bve.gen1,
              path = "maternal_plink", fam.id = TRUE, type = 2)
    
    write.table(file = "EBV.txt", cbind(get.id(population, gen = bve.gen1), ebv), row.names = FALSE, col.names = FALSE, quote = FALSE)
    write.table(file = "sol.dat", cbind(1,1,1,1:length(alpha_hat), alpha_hat), row.names = FALSE, col.names = FALSE, quote = FALSE)
    write.table(file = "all_freq", cbind(1:length(alpha_hat), 0), row.names = FALSE, col.names = FALSE, quote = FALSE)
    
    system("./msv --matgeno maternal_plink.bed --patgeno paternal_plink.bed --freq all_freq --ebv EBV.txt --sol sol.dat --snpeff 1 --nrgen 2 --selprop 0.1 --bp2M 1e-08")
    
    a = read.table("EBV.txt")
    b = read.table("genetic_MSVs_1gen.txt")
    c = read.table("Index_5_6.txt") #Piter Index 5/6
    d = read.table("ExpBVSelGrOff_per_individual.txt") # Tobias Index
    
    write.table(file = "sol.dat", cbind(1,1,1,1:length(alpha_real), alpha_real), row.names = FALSE, col.names = FALSE, quote = FALSE)
    
    system("./msv --matgeno maternal_plink.bed --patgeno paternal_plink.bed --freq all_freq --ebv EBV.txt --sol sol.dat --snpeff 1 --nrgen 1 --selprop 0.1 --bp2M 1e-08")
    a1 = read.table("EBV.txt")
    b1 = read.table("genetic_MSVs_1gen.txt")
    c1 = read.table("Index_5_6.txt") #Piter Index 5/6
    d1 = read.table("ExpBVSelGrOff_per_individual.txt") # Tobias Index
    
    
    new_ebv = d[,2]
    
    insert.bve = cbind(names(ebv), new_ebv)
    population = insert.bve(population, bves = insert.bve )
    
    acc3 = c(acc3,     cor(b[,2],b1[,2]))
    
    acc2 = rbind(acc2, t(analyze.bv(population, gen = max(bve.gen))[[1]]))
    
    pop1 = breeding.diploid(population, bve = TRUE, bve.gen = bve.gen)
    
    acc1 = rbind(acc1, t(analyze.bv(pop1, gen = max(bve.gen))[[1]]))
    
    print(colMeans(acc1))
    print(colMeans(acc2))
    
    # Selection based on genomic breeding value
    population = breeding.diploid(population, selection.size = sel_size, breeding.size = pop_size,
                                  selection.criteria = "bve",
                                  share.genotyped = 1,
                                  display.progress = display.progress)
    
    
    
  } else if( scenario == 336){
    
    
    work_dir = paste0("mixblup_", scenario, "_", seed)
    if(index == 1){
      dir.create(work_dir)
      system(paste0("cp /home/WUR/pook001/SysDir.inp ", work_dir, "/SysDir.inp"))
      setwd(work_dir)
      
      system(paste0("cp /home/WUR/pook001/msv msv"))
      system("chmod +x msv")
    }
    
    get.plink(population, gen =bve.gen1,
              path = "paternal_plink", fam.id = TRUE, type = 1)
    
    get.plink(population, gen =bve.gen1,
              path = "maternal_plink", fam.id = TRUE, type = 2)
    
    write.table(file = "EBV.txt", cbind(get.id(population, gen = bve.gen1), ebv), row.names = FALSE, col.names = FALSE, quote = FALSE)
    write.table(file = "sol.dat", cbind(1,1,1,1:length(alpha_hat), alpha_hat), row.names = FALSE, col.names = FALSE, quote = FALSE)
    write.table(file = "all_freq", cbind(1:length(alpha_hat), 0), row.names = FALSE, col.names = FALSE, quote = FALSE)
    
    system("./msv --matgeno maternal_plink.bed --patgeno paternal_plink.bed --freq all_freq --ebv EBV.txt --sol sol.dat --snpeff 1 --nrgen 1 --selprop 0.1 --bp2M 1e-08")
    
    a = read.table("EBV.txt")
    b = read.table("genetic_MSVs_1gen.txt")
    c = read.table("Index_5_6.txt") #Piter Index 5/6
    #d = read.table("ExpBVSelGrOff_per_individual.txt") # Tobias Index
    
    write.table(file = "sol.dat", cbind(1,1,1,1:length(alpha_real), alpha_real), row.names = FALSE, col.names = FALSE, quote = FALSE)
    
    system("./msv --matgeno maternal_plink.bed --patgeno paternal_plink.bed --freq all_freq --ebv EBV.txt --sol sol.dat --snpeff 1 --nrgen 1 --selprop 0.1 --bp2M 1e-08")
    a1 = read.table("EBV.txt")
    b1 = read.table("genetic_MSVs_1gen.txt")
    c1 = read.table("Index_5_6.txt") #Piter Index 5/6
    #d1 = read.table("ExpBVSelGrOff_per_individual.txt") # Tobias Index
    
    
    new_ebv = c[,2]
    
    insert.bve = cbind(names(ebv), new_ebv)
    population = insert.bve(population, bves = insert.bve )
    
    acc3 = c(acc3,     cor(b[,2],b1[,2]))
    
    acc2 = rbind(acc2, t(analyze.bv(population, gen = bve.gen)[[1]]))
    
    pop1 = breeding.diploid(population, bve = TRUE, bve.gen = bve.gen)
    
    acc1 = rbind(acc1, t(analyze.bv(pop1, gen = bve.gen)[[1]]))
    
    print(colMeans(acc1))
    print(colMeans(acc2))
    
    # Selection based on genomic breeding value
    population = breeding.diploid(population, selection.size = sel_size, breeding.size = pop_size,
                                  selection.criteria = "bve",
                                  share.genotyped = 1,
                                  display.progress = display.progress)
    
    
    
  } else if( scenario == 337){
    
    
    work_dir = paste0("mixblup_", scenario, "_", seed)
    if(index == 1){
      dir.create(work_dir)
      system(paste0("cp /home/WUR/pook001/SysDir.inp ", work_dir, "/SysDir.inp"))
      setwd(work_dir)
      
      system(paste0("cp /home/WUR/pook001/msv msv"))
      system("chmod +x msv")
    }
    
    get.plink(population, gen =bve.gen1,
              path = "paternal_plink", fam.id = TRUE, type = 1)
    
    get.plink(population, gen =bve.gen1,
              path = "maternal_plink", fam.id = TRUE, type = 2)
    
    write.table(file = "EBV.txt", cbind(get.id(population, gen = bve.gen1), ebv), row.names = FALSE, col.names = FALSE, quote = FALSE)
    write.table(file = "sol.dat", cbind(1,1,1,1:length(alpha_hat), alpha_hat), row.names = FALSE, col.names = FALSE, quote = FALSE)
    write.table(file = "all_freq", cbind(1:length(alpha_hat), 0), row.names = FALSE, col.names = FALSE, quote = FALSE)
    
    system("./msv --matgeno maternal_plink.bed --patgeno paternal_plink.bed --freq all_freq --ebv EBV.txt --sol sol.dat --snpeff 1 --nrgen 1 --selprop 0.1 --bp2M 1e-08")
    
    a = read.table("EBV.txt")
    b = read.table("genetic_MSVs_1gen.txt")
    c = read.table("Index_5_6.txt") #Piter Index 5/6
    #d = read.table("ExpBVSelGrOff_per_individual.txt") # Tobias Index
    
    write.table(file = "sol.dat", cbind(1,1,1,1:length(alpha_real), alpha_real), row.names = FALSE, col.names = FALSE, quote = FALSE)
    
    system("./msv --matgeno maternal_plink.bed --patgeno paternal_plink.bed --freq all_freq --ebv EBV.txt --sol sol.dat --snpeff 1 --nrgen 1 --selprop 0.1 --bp2M 1e-08")
    a1 = read.table("EBV.txt")
    b1 = read.table("genetic_MSVs_1gen.txt")
    c1 = read.table("Index_5_6.txt") #Piter Index 5/6
    #d1 = read.table("ExpBVSelGrOff_per_individual.txt") # Tobias Index
    
    
    new_ebv = c[,3]
    
    insert.bve = cbind(names(ebv), new_ebv)
    population = insert.bve(population, bves = insert.bve )
    
    acc3 = c(acc3,     cor(b[,2],b1[,2]))
    
    acc2 = rbind(acc2, t(analyze.bv(population, gen = bve.gen)[[1]]))
    
    pop1 = breeding.diploid(population, bve = TRUE, bve.gen = bve.gen)
    
    acc1 = rbind(acc1, t(analyze.bv(pop1, gen = bve.gen)[[1]]))
    
    print(colMeans(acc1))
    print(colMeans(acc2))
    
    # Selection based on genomic breeding value
    population = breeding.diploid(population, selection.size = sel_size, breeding.size = pop_size,
                                  selection.criteria = "bve",
                                  share.genotyped = 1,
                                  display.progress = display.progress)
    
    
    
  } else if( scenario == 338){
    
    acc2 = rbind(acc2, t(analyze.bv(population, gen = max(bve.gen))[[1]]))
    
    # Selection based on genomic breeding value
    population = breeding.diploid(population, selection.size = sel_size, breeding.size = pop_size,
                                  selection.criteria = "bve",
                                  share.genotyped = 1,
                                  display.progress = display.progress)
    
  } else if(scenario == 396){
    
    
    sel_size_tmp = sel_size
    # selection on the female side is done without OGC, on male male size selection.size = population size
    # to provide optiSel with all potential sires
    # calculation on how to set ub.sKin can be tricky has this corresponds to the increase of inbreeding
    # that is caused by the male side of the breeding program
    sel_size_tmp[1] = pop_size[1]
    # Selection based on genomic breeding value
    population = breeding.diploid(population, selection.size = sel_size_tmp,
                                  breeding.size = pop_size,
                                  selection.criteria = "bve",
                                  share.genotyped = 1,
                                  ogc = TRUE,
                                  ogc.target = "max.BV",
                                  ogc.uniform = "female",
                                  ogc.ub.sKin.increase = 0.0043,
                                  display.progress = display.progress)
    
  } else if(scenario == 397){
    
    
    sel_size_tmp = sel_size
    # selection on the female side is done without OGC, on male male size selection.size = population size
    # to provide optiSel with all potential sires
    # calculation on how to set ub.sKin can be tricky has this corresponds to the increase of inbreeding
    # that is caused by the male side of the breeding program
    sel_size_tmp[1] = pop_size[1]
    # Selection based on genomic breeding value
    population = breeding.diploid(population, selection.size = sel_size_tmp,
                                  breeding.size = pop_size,
                                  selection.criteria = "bve",
                                  share.genotyped = 1,
                                  ogc = TRUE,
                                  ogc.target = "max.BV",
                                  ogc.uniform = "female",
                                  ogc.ub.sKin.increase = 0.003,
                                  display.progress = display.progress)
    
  } else if(scenario == 398){
    
    
    sel_size_tmp = sel_size
    # selection on the female side is done without OGC, on male male size selection.size = population size
    # to provide optiSel with all potential sires
    # calculation on how to set ub.sKin can be tricky has this corresponds to the increase of inbreeding
    # that is caused by the male side of the breeding program
    sel_size_tmp[1] = pop_size[1]
    # Selection based on genomic breeding value
    population = breeding.diploid(population, selection.size = sel_size_tmp,
                                  breeding.size = pop_size,
                                  selection.criteria = "bve",
                                  share.genotyped = 1,
                                  ogc = TRUE,
                                  ogc.target = "max.BV",
                                  ogc.uniform = "female",
                                  ogc.ub.sKin.increase = 0.004,
                                  display.progress = display.progress)
    
  } else if(scenario == 399){
    
    
    sel_size_tmp = sel_size
    # selection on the female side is done without OGC, on male male size selection.size = population size
    # to provide optiSel with all potential sires
    # calculation on how to set ub.sKin can be tricky has this corresponds to the increase of inbreeding
    # that is caused by the male side of the breeding program
    sel_size_tmp[1] = pop_size[1]
    # Selection based on genomic breeding value
    population = breeding.diploid(population, selection.size = sel_size_tmp,
                                  breeding.size = pop_size,
                                  selection.criteria = "bve",
                                  share.genotyped = 1,
                                  ogc = TRUE,
                                  ogc.target = "max.BV",
                                  ogc.uniform = "female",
                                  ogc.ub.sKin.increase = 0.006,
                                  display.progress = display.progress)
    
  } else if(scenario == 400){
    
    
    sel_size_tmp = sel_size
    # selection on the female side is done without OGC, on male male size selection.size = population size
    # to provide optiSel with all potential sires
    # calculation on how to set ub.sKin can be tricky has this corresponds to the increase of inbreeding
    # that is caused by the male side of the breeding program
    sel_size_tmp[1] = pop_size[1]
    # Selection based on genomic breeding value
    population = breeding.diploid(population, selection.size = sel_size_tmp,
                                  breeding.size = pop_size,
                                  selection.criteria = "bve",
                                  share.genotyped = 1,
                                  ogc = TRUE,
                                  ogc.target = "max.BV",
                                  ogc.uniform = "female",
                                  ogc.ub.sKin.increase = 0.007,
                                  display.progress = display.progress)
    
  } 
  
  
  
}

# Generate offspring from the best animals based on traditional EBVs
for(index in 1:cycles){
  print(index)
  current_gen = get.ngen(population) - cycles
  
  sel_crit = "bve"
  if(scenario %in% c(320:326)){
    sel_crit = "bv"
  }
  
  bve.gen = current_gen
  if(scenario %in% 335:338){
    bve.gen = (current_gen-5):current_gen
  }
  population = breeding.diploid(population, bve = TRUE, bve.gen = bve.gen)
  
  population = breeding.diploid(population, selection.size = sel_size, breeding.size = pop_size,
                                selection.m.database = cbind(current_gen,1),
                                selection.f.database = cbind(current_gen,2),
                                selection.criteria = sel_crit,
                                share.genotyped = 1,
                                display.progress = display.progress)
}


#### Evaluation of the population


#### Evaluation of the population
ngen = get.ngen(population)

bv = numeric(ngen)
bv_sd = numeric(ngen)
inb = numeric(ngen)
share_fixed = numeric(ngen)
share_fixed_qtl = numeric(ngen)
share_fixed_good = numeric(ngen)
share_fixed_bad = numeric(ngen)
for(index in 1:ngen){
  print(index)
  bv[index] = mean(get.bv(population, gen = index))
  bv_sd[index] = sd(get.bv(population, gen = index))
  inb[index] = mean(inbreeding.emp(population, gen = index))
  
  geno = get.geno(population, gen = index)
  p = rowMeans(geno)/2
  share_fixed[index] = mean( p ==0 | p ==1)
  share_fixed_qtl[index] = mean( (p ==0 | p ==1)[population$info$effect.p])
  
  share_fixed_good[index] = mean(((alpha_real > 0 & p ==1) | (alpha_real < 0 & p ==0))[population$info$effect.p])
  share_fixed_bad[index] = mean(((alpha_real < 0 & p ==1) | (alpha_real > 0 & p ==0))[population$info$effect.p])
}


save(file = title,
     list = c("bv", "bv_sd", "inb", "share_fixed", "share_fixed_qtl", "share_fixed_good", "share_fixed_bad"))
```
