## Supplementary Files 15 &16 for "Strategies to improve selection compared to selection based on estimated breeding values": Supplementary_FileS16.R

```
#########################################################################################################
##################################### Plot script for manuscript: #######################################
#######  Strategies to improve selection  compared to selection based on estimated breeding values ######
#########################################################################################################

# agreegating results from simulations
n = 50 
nsc = 400

bv_total = bv_sd_total = inb_total= share_fixed_total = share_fixed_qtl_total =
  share_fixed_good_total = share_fixed_bad_total = matrix(0, nrow=nsc, ncol=111)
for(scenario in (1:nsc)){

    print(scenario)
  n_temp = 0
  scenario_tmp = scenario
  
  # scenarios that used 500 replicates
  
  if(scenario %in% c(187,189 ,190,188, 191, 192, 193, 299)){
    n = 500
    if(scenario==299){
      scenario_tmp = 1
    }
  } else{
    n = 50
  }
  for(seed in c(1:n)){
    
    # In the final results presented so simulations failed. This is just a safety net when analyzing semi-finished scenarios
    tryCatch(  {
    
      load(paste0("uniqueness_v3/sc", scenario_tmp , "seed", seed, "_potential.RData"))

      bv_total[scenario,] = bv_total[scenario,] + bv
      bv_sd_total[scenario,] = bv_sd_total[scenario,] + bv_sd
      inb_total[scenario,] = inb_total[scenario,] + inb
      share_fixed_total[scenario,] = share_fixed_total[scenario,] + share_fixed
      share_fixed_qtl_total[scenario,] = share_fixed_qtl_total[scenario,] + share_fixed_qtl
      share_fixed_good_total[scenario,] = share_fixed_good_total[scenario,] + share_fixed_good
      share_fixed_bad_total[scenario,] = share_fixed_bad_total[scenario,] + share_fixed_bad
      
      n_temp = n_temp + 1
    },
    error = function(e) {})
  }
  
  bv_total[scenario,] = bv_total[scenario,]/n_temp
  bv_sd_total[scenario,] = bv_sd_total[scenario,]/n_temp
  inb_total[scenario,] = inb_total[scenario,]/n_temp
  share_fixed_total[scenario,] = share_fixed_total[scenario,]/n_temp
  share_fixed_qtl_total[scenario,] = share_fixed_qtl_total[scenario,] /n_temp
  share_fixed_good_total[scenario,] = share_fixed_good_total[scenario,]/n_temp
  share_fixed_bad_total[scenario,] = share_fixed_bad_total[scenario,]/n_temp
  
  print(n_temp)
}


## Plot of a single scenario

{
  png("C:/Users/pook001/OneDrive - Wageningen University & Research/Figure_2_unique.png",
      width = 4000, height = 1300, res = 300)
  
  index = 1
  limits = c(-0.2,1.5)
  
  par(mfrow=c(1,4))
  plot(0:50,bv_total[index,c(11,62:111)], xlim=c(0,50), 
       type = "l", lwd = 2,
       main = "genetic gain", ylab = "genetic gain in gSD", xlab="generation")
  
  coords <- par("usr")
  text(0.5,0.98 * coords[4] + 0.02 * coords[3], "a")
  
  plot(0:50, inb_total[index,11:61], xlim=c(0,50), lwd = 2, type = "l", ylim=c(0,1), main = "inbreeding", ylab = "inbreeding level", xlab="generation")
  
  par(new = TRUE)  # Overlay a new plot
  
  plot(1:50, diff(inb_total[index,11:61]), lwd = 2, type ="l", col = 2, axes = FALSE, xlab = "", ylab = "")
  axis(side = 4, col = 2, col.axis = 2, las = 3)  # Add right y-axis
  mtext("inbreeding rate", side = 4, line = 1.6, col = 2, cex = 0.7)  # Label for right y-axis
  coords <- par("usr")
  text(0.5,0.98 * coords[4] + 0.02 * coords[3], "b")
  
  plot(0:50, bv_sd_total[index,11:61], type = "l", lwd =2,  ylim=c(0,1), main = "genetic variance", ylab = "genetic variance", xlab="generation")
  coords <- par("usr")
  text(0.5,0.98 * coords[4] + 0.02 * coords[3], "c")
  
  plot(0:50, share_fixed_qtl_total[index,11:61], type = "l", lwd = 2, xlim=c(0,50), ylim=c(0,1), main = "Fixed QTLs", ylab = "share", xlab="generation")
  
  lines(0:50, share_fixed_total[index,11:61], type = "l", lwd = 2, col = 4)
  
  lines(0:50, share_fixed_good_total[index,11:61] / share_fixed_qtl_total[index,11:61], type = "l", lwd = 2, col = 2)
  
  coords <- par("usr")
  text(0.5,0.98 * coords[4] + 0.02 * coords[3], "d")
  
  legend("topright", c("QTLs fixed", "SNPs fixed", "Share QTLs fixed benefical"), 
         col = c(1,4,2), lty = 1, lwd =3, cex =0.8)
  
  
  dev.off()
}

## Plot of multiple scenario against the reference

{
  
  png("C:/Users/pook001/OneDrive - Wageningen University & Research/Figure_5_unique.png",
      width = 2600, height = 1600, res = 300)
  legend = NULL
  
  legend[c(1, 95, 96, 97,98,99,100,
           118, 119, 120,121
  )] = c("EBVs","no fullsib-matings",
         "no halfsib-matings",
         "inbreeding quantile: 0.90",
         "inbreeding quantile: 0.50",
         "inbreeding quantile: 0.25",
         "inbreeding quantile: 0.10",
         "kinship quantile: 0.90",
         "kinship quantile: 0.50",
         "kinship quantile: 0.25",
         "kinship quantile: 0.10"
  )
  
  limits = c(-0.7,1.4)
  used_color = NULL
  par(mfrow=c(1,2))
  
  # 67, 174
  include = c( 95, 96, 97,98,99,100,  118,119,120,121)
  
  used_color[include] = 1:8
  used_color[120:121] = c("brown", "darkgreen")
  plot(-100,0, xlim=c(0,50), ylim=limits, main = "genetic gain (relative to baseline)", ylab = "genetic gain in gSD", xlab="generation")
  for(index in include){
    #  lines(0:50, bv_total[index,11:61] - bv_total[1,11:61], col=used_color[index], lwd=2)
    lines(0:50, bv_total[index,c(11,62:111)] - bv_total[1,c(11,62:111)], lty = 1, col=used_color[index], lwd=2)
    
  }
  abline(h = 0, lty = 2)
  
  text(0,1.4, "a")
  legend("top", as.character(legend[include]),
         col=used_color[include], lty = 1, lwd=2, cex = 0.6)
  
  
  plot(-10000,0, xlim=c(0,50), ylim=c(-0.11,0.10), main = "inbreeding (relative to baseline)", ylab = "inbreeding level", xlab="generation")
  for(index in include){
    lines(0:50,  (inb_total[index,11:61] - inb_total[1,11:61]) , col=used_color[index], lwd=2)
  }
  text(0,0.1, "b")
  
  abline(h = 0, lty = 2)
  dev.off()dev.off()dev.off()
  
  index = 98
  bv_total[index,c(11,62:111)] - bv_total[1,c(11,62:111)]
  (bv_total[index,c(11,62:111)] - bv_total[1,c(11,62:111)]) / (bv_total[1,c(11,62:111)] -bv_total[1,c(11)] )
  
  (inb_total[index,11:61] - inb_total[1,11:61])
  (inb_total[index,11:61] - inb_total[1,11:61]) /   (inb_total[1,11:61] - inb_total[1,11])
}
```
